# An Efficient Computing Theory of Prefrontal Structured Working Memory Representations

**DOI:** 10.64898/2026.02.16.706126

**Authors:** William Dorrell, Peter E. Latham, Timothy E. J. Behrens, James C. R. Whittington

## Abstract

The efficient coding hypothesis presents a compelling success story for theoretical and systems neuroscience. It marshals a unifying idea, that neural codes can be understood as efficient encodings of natural stimuli, to explain phenomena from across sensory systems, sometimes with exquisite precision. However, similar normative assaults on cognitive representations, such as those in prefrontal or entorhinal cortex, have been less comprehensively successful. We argue that this reflects efficient coding’s focus on encoded variables, overlooking the computations that neural circuits implement. Here, instead, we develop an efficient *computing* theory that studies optimal implementations of recurrent cognitive computations. We apply this framework to the prefrontal cortex, in particular, to a rich vein of neural recordings: structured working memory tasks, such as recalling a sequence. Despite using near-identical tasks, the literature has reported two distinct coding schemes: one contextual, with neurons active only in a particular sequence, the other compositional, with neurons tuned to a single sequence element. Just as efficient coding links stimuli statistics to optimal representation, our theory relates task structure and statistics to optimal representation. In so doing, we find that compositional and contextual codes can be understood as two extremes of a spectrum of optimal representations, with the correlations amongst sequence elements determining a task’s position on this spectrum. Our theory highlights previously underappreciated discrepancies between measured representations, explaining them via subtle task differences; and allows us to infer the algorithm underlying otherwise ambiguous neural data. In sum, we demonstrate an efficient computing approach that makes normative statements about cognitive representations, and serves as a tool for understanding a swathe of neural data.

## 1 Introduction

The efficient coding hypothesis is one of the most successful and longstanding theories in neuroscience. In short, it argues that neural activity can be understood as the minimal-energy, faithful encoding of relevant variables (Attneave; Barlow). This principle, stunning in its simplicity, has found widespread use across early sensory processing, explaining the relationship between nonlinearities and stimulus distribution (Laughlin), the distribution of retinal ON and OFF cells (Ratliff et al.), the alignment of retinal lattices (Jun, Field, and Pearson), the structure of the retinal ganglion cell (Atick and Redlich), V1 simple cell (Olshausen and Field), or cochlear representations (Smith and Lewicki), and much else besides. As a theory of early sensory representations, efficient coding is well-framed and useful.

However, there have been fewer successful applications of efficient coding to cognitive representations. Existing cortical applications of efficient coding have largely been sensory (Coen-Cagli, Kohn, and Schwartz; Młynarski and Tkačik; Hermundstad et al.; Tkačik et al.) and are significantly less comprehensive (Carandini et al.; Olshausen and Field; Stringer et al.). We argue this reflects a mismatch in assumptions. An efficient code is optimised for communication: it compactly summarises the required information permitting low cost transmission. This aligns with the sensory periphery, for example, retinal ganglion cells, which transmit retinal representations for further cortical processing via a tight bottleneck, the optic nerve. But cognitive representations are not just for communicating; instead, they form the substrate for the brain’s ongoing computations. We should therefore expect these representations, and our theories of them, to reflect the relevant computations.

In this work we therefore study an efficient *computing* theory, which models neural activity as the most efficient implementation of a given algorithm. This combines efficient coding arguments with constraints that ensure the population can perform the posited computation. We have previously used these approaches to model the grid cell representation as the optimal implementation of path-integration (Dorrell et al.; Dorrell and Whittington). Here, we turn to prefrontal cortex.

The prefrontal cortex is hypothesised to form the mind’s workspace, encoding relevant stimuli and using them to construct flexible behavioural strategies. Responding flexibly requires, among other things, first, using context to separate otherwise identical scenarios, and, second, composing an appropriate response from previously learnt primitives. For example, the sensory tableau upon entering meetings with your advisor might be identical, but contextual information about their recent Nobel prize/firing for corruption of the youth (delete as appropriate) will modify the scene’s meaning. Faced with these different scenarios, appropriate behaviour then requires composing congratulation/condemnation with a kindly worded request for that delayed reference letter.

Prefrontal neural recordings show evidence of both of these kinds of flexibility. On the one hand, Xie et al. trained monkeys to saccade to a sequence of remembered location. During the delay period of this task they found that neural activity was composed of three subspaces, the projection of the activity into each subspace encoded one of the remembered locations, fig. 1A. This representation is overtly compositional, since it is the sum of three parts, each of which encodes a single memory. On the other hand, Shima and Tanji trained monkeys to perform a set of three remembered movements in different orders. They found that single neurons fired in response to only one order, fig. 1B, (Shima and Tanji). We will refer to this as contextual coding, in which, analogously to splitter cells whose firing distinguishes the same position in different contexts (Frank, Brown, and Wilson; Wood et al.), single neurons encode ‘beginnning of pull-turn-push’ sequence, rather than encoding a single upcoming action, fig. 1B. Findings of these two types of code, illustrated in fig. 1C, are relatively general, with many works reporting prefrontal recordings that can be seen as either contextual (Lu, Matsuzawa, and Hikosaka; Takahashi et al.; Ma, Zhang, and Zhou; Schuck et al.; Zhou et al.) or compositional (El-Gaby et al.; Basu et al.; Mushiake et al.; Panichello and Buschman), and RNNs trained on similar tasks have recovered the observed compositional coding (Botvinick and Plaut; Piwek, Stokes, and Summerfield; Whittington et al.; Wang, Fusi, and Stachenfeld).

**Figure 1.**
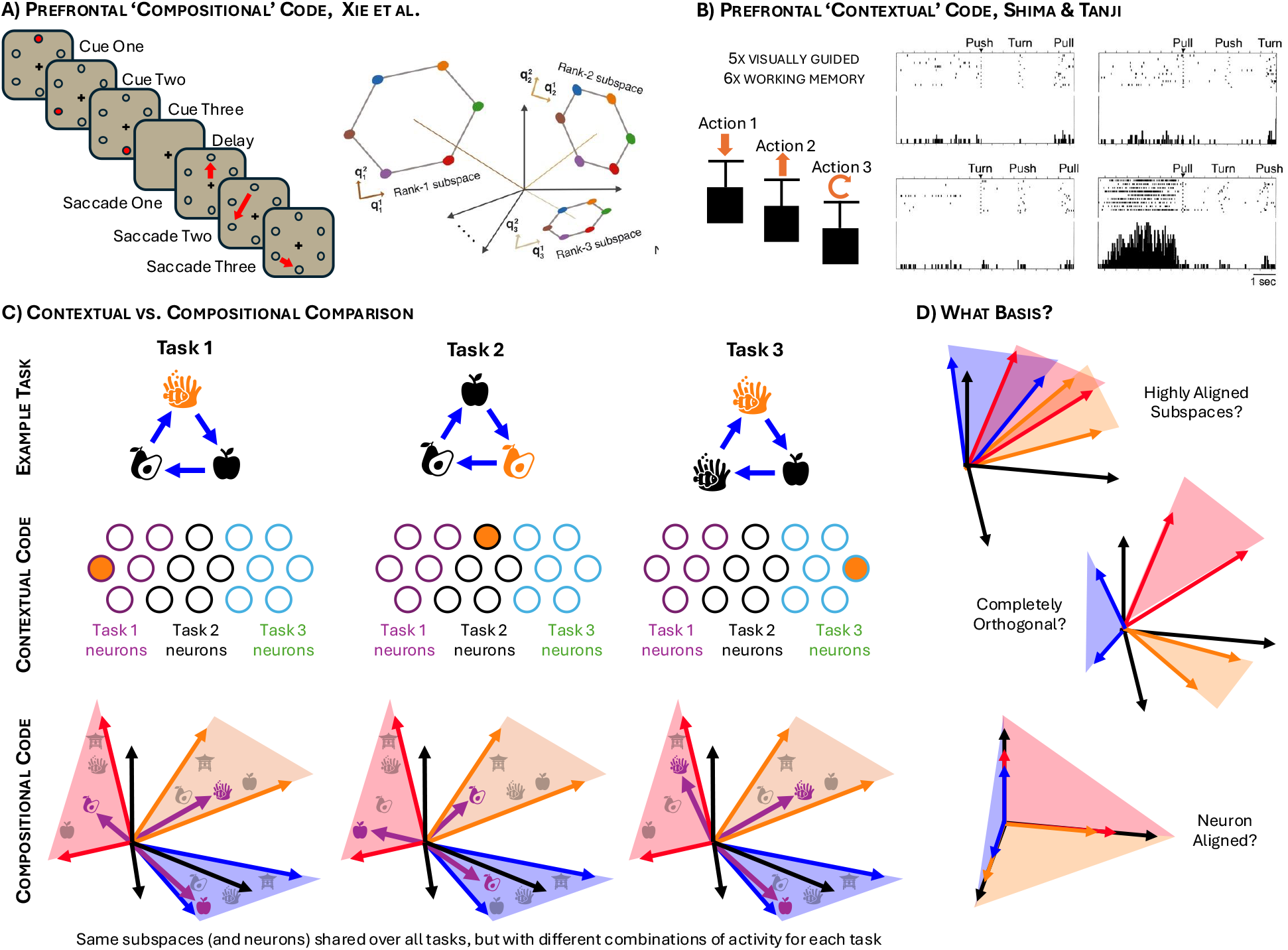
Contextual and compositional codes are found in prefrontal cortex. **A)** Left, Xie et al. trained monkeys to observe then saccade to a sequence of presented cues. Right, the measured neural activity was well described as the sum of three subspaces each encoding a sequence element. **B)** Left, Shima & Tanji trained monkeys to perform remembered sequences of movements. Right, They found neurons in the pre-SMA that fired in preparation for only one sequence. **C)** Contextual and compositional codes seem markedly different. Consider a family of three tasks, each a simple loop of 3 stimuli. A contextual code would use different neurons (subspaces) for each task (here neurons represent positions in each task), middle row, while a compositional code shares neurons (subspaces) across tasks and encodes each task using different compositions of projections within the subspaces (purple arrows show the activity vector in each subspace), bottom row. We find that these contextual and compositional codes can be seen as extremes of a spectrum of subspace-based codes. Hence, **(D)** a key question we consider is how to optimally arrange subspaces, both with respect to one another, and to the neuron basis.

This diversity of recordings is puzzling, especially from identical tasks like reporting a remembered sequence (Shima and Tanji; Xie et al.). To understand this diversity we introduce an efficient computing theory of structured working memory representations. We consider representations that are able to recall the correct item according to the underlying structural rules of the task. Subject to this computational constraint, we study the most energy efficient representation.

We find that the optimal solution depends heavily on the structure and statistics of the task. When the different elements of a structured working memory task are independent, the optimal representation is compositional, using an orthogonal subspace per element. Conversely, when elements are highly correlated, they are encoded in aligned subspaces, which manifest as contextually-tuned single neurons. This unifies both representations as extremes on an axis of task diversity. Further, the theory makes precise predictions about the alignment and sizes of the encoding subspaces, and we find that our predictions can explain underappreciated discrepancies between measured recordings via subtle differences in the task. Finally, in representations consistent with multiple subspace algorithms (how neural activity flows between subspaces during a task), we find that only one algorithm would be efficiently implemented in the manner observed, making a clear prediction for future experiments.

In sum, we develop an efficient computing approach that applies normative efficient-coding style reasoning to cognitive representations, and find that it can explain and predict detailed aspects of prefrontal working memory representations.

### 2 An Efficient Computing Framework for Structured Working Memory Tasks

In this section we outline a family of structured working memory tasks, designed to encompass many of those studied. We then outline our efficient computing model of the optimal representation within these tasks.

### 2.1 Structured working memory tasks

Each of our tasks comprises three parts: (1) latent states, (2) actions that transition the agent between latent states, and (3) stimuli attached to latent states. For example, the latent states might be positions in a room, the actions the movements between them (like ‘step north’), and the stimuli could be an object at each point in the room, fig. 2A. The agent experiences many environments that share the same structure, such as 2D space, but with a different pairing of stimuli to state in each environment. The agent is told the particular stimuli-state pairing in the current environment, and a sequence of actions, and has to recall the stimuli associated to each latent state it visits in sequence, fig. 2D.

**Figure 2.**
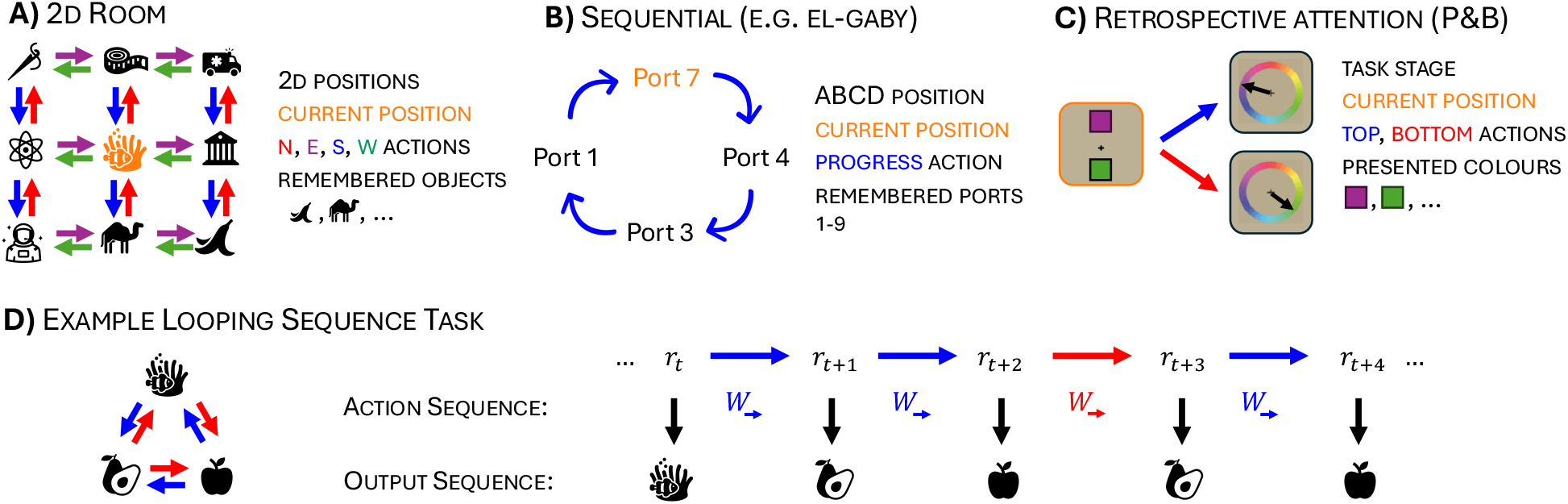
Structured working memory tasks and our theoretical formalism. **A)** An example 2D room task: states are positions in the room, there is an object at each position, and the agent moves using four actions, N, S, E, W. **B)** An example looping sequential task, as in El-Gaby et al.: states are positions in the loop, the only action is progress forward, and items are the remembered rewarded port locations. **C)** An example retrospective attention task, as in Panichello and Buschman: in the first state the presented item includes the two colours. Depending on the subsequent actions, top or bottom colour, the agent has to recall the corresponding colour. **D)** To understand the optimal representation, ***r***_*t*_, of structured memory tasks, we summarise each task as comprising a sequences of actions 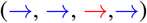 and corresponding stimuli that must be predicted (recalled). Our theoretical formalism then asks what the representation, ***r***_*t*_, must look like—after training on a family of such tasks with shared structure (e.g. 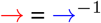) but different stimuli—in order to successfully predict (recall) each observation after each action.

A simple example, to which we will return, is sequence working memory (Shima and Tanji; Xie et al.; Mushiake et al.; Basu et al.; El-Gaby et al.). For example, El-Gaby et al. train mice in an arena with 9 ports, of which four are rewarded in a particular sequence: an ‘ABCD’ task. Example port sequences are 74317431 …, or 81978197 …, and so on. Encoded in our framework, the latent state would be sequence element (A, B, C, or D), there would be only one action ‘progress forward’ that would map A →B →C →D →A, and the latent-stimuli pairing would describe which of the 9 possible ports was mapped to each of the 4 latent states, fig. 2B (e.g. if {*A, B, C, D*} = {7, 4, 3, 1} then 74317431… is correct). An example with more than one action is the retrospective attention task of Panichello and Buschman in which, similarly, a set of cues are presented, in this case two—one above the other. Two actions then correspond to selecting one of the cues to report, top or bottom, fig. 2C.

### 2.2 The Efficient Computing Hypothesis

In this work we build a model of the optimal neural representation, ***r***(*s, e*), a vector of neural firing rates that is a function of both the agent’s current latent state, *s*, and environment, *e*. Like the efficient coding hypothesis, our theory takes the form of an optimisation problem, whose solution is our model of neural activity. Conventional efficient coding approaches contain at least two parts: one functional that asks the representation to meaningfully encode the relevant variables, and a second that enforces biological plausibility/efficiency. For example, Olshausen and Field study a nonnegative sparse visual representation (biological considerations) that is optimised to allow linear reconstruction of images (functional consideration). Our efficient computing approach supplements these biological and functional components with a third ‘computing constraint’ that ensures the representation is able to ‘navigate’ the latent states, *s*, and thus subserve the relevant algorithm.

Concretely, we optimise each vector ***r***(*s, e*), along with a series of related weight matrices, such that an efficiency loss is minimised while satisfying the relevant algorithmic constraints, which we detail below. We do this simultaneously for all latent states, *s*, and environments, *e*, in our training dataset. The constraints make the optimal representations highly structured, ensuring it can be used to solve our structured working memory tasks in a prescribed way.

#### Functional Constraint

First the representation must be able to recall the stimuli associated to the current state, which we enforce with an affine decoder that is fixed across all states and environments:

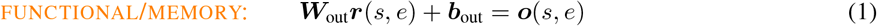

where ***o***(*s, e*) is the label corresponding to the stimuli associated to state *s* in environment *e*. This equation says that we can use the representation to easily predict the relevant stimuli, for example, as the agent traverses the 2D room example, fig. 2A, there will be different representations for each *s* but that a fixed decoder, ***W***_out_, will be able to decode the relevant stimuli at each state, *s*, for each environment, *e*.

#### Computing Constraint

Second, our ‘computing constraint’ forces the representation to perform the required computation. Here, this means that as the agent moves from latent state to latent state, e.g. moving around a room, fig. 2A, or progressing through a task, fig. 2B-C, the representation must update appropriately depending on the action taken. We enact this similarly with a set of affine transformations, one for each action:

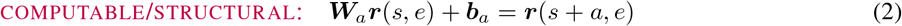

This equation ensures that the prefrontal internal model accurately captures the rules of the external world, allowing the animal to navigate internally through the task and predict the corresponding stimuli. For example, if there’s a conceptual action, like ‘step east’, that links together many states, fig. 2A, there is a corresponding neural transformation, ***W***_east_, that does the same to the representation. In short, it is eq. (2) that makes the representation an internal model.

The combination of eq. (1) and eq. (2) ensures the representation can solve the task. To see this we can follow the logic of an update: after taking an action the agent updates its representation, eq. (2), and can then decode the stimuli relevant to the new state: ***W***_out_***r***(*s* + *a, e*) + ***b***_out_ = ***o***(*s* + *a, e*). Further, crucially, the transformations, ***W***_out_ and ***W***_*a*_, are identical across state and environment. This ensures that, once you have learnt the rules of the current environments (e.g. 2D space, or loops), you could use the same set of transformations to solve a previously-unseen sequence.

#### Biological Constraint

Subject to these task-solving constraints, we study the most energy efficient representation. We do this by minimising the following energy-use loss, which sums the energy used in neural activity and in weights, matching evidence that a significant part of spiking energy-use is in synaptic transmission (Harris, Jolivet, and Attwell)^2^:

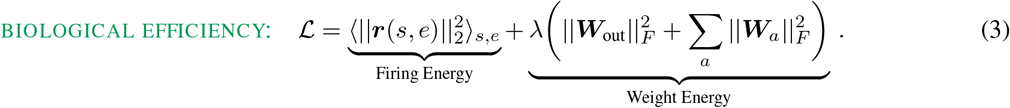

Minimising eq. (3) subject to eq. (1) and eq. (2) comprises the simplest version of our efficient computing theory. To do this, we not only optimise ***r***(*s, e*), but also ***W***_out_ and the set of action matrices, ***W***_*a*_.

This formalises our structured working memory task family, and our efficient computing optimisation problem. We lightly adapt our framework depending on the situation, appendix D, in particular, we sometimes include an input matrix, and when modelling single neurons we constrain the neural activity to be non-negative.

Our analysis now proceeds in two parts. First, we do mathematical analysis to extract a general conclusion: solving structured working memory tasks with (gated) linear recurrent weights requires the representation to construct a series of subspaces. Then we numerically optimise the representations and link the resulting neural patterns to task structure and statistics.

## 3 From Compositional to Contextual: A Spectrum of Subspace Solutions

In this section we intuitively describe how the constraints of solving the task, eq. (1) and eq. (2), imply that the representation is composed of a set of neural subspaces, each encoding a different task element (Details: Appendix B.1). Despite this universally ‘compositional’ structure, minimising energy use, eq. (3), creates large differences between the encodings of different tasks, leading to representations that appear either compositional or contextual depending on the task statistics. In particular, highly correlated memories are optimally encoded in aligned subspaces, while independent memories are encoded in orthogonal subspaces. These two extremes sketch an axis of task diversity onto which each experiment can be placed, allowing us to relate different task statistics to the measured discrepancies in neural subspace alignments.

### 3.1 Solving Structured Working Memory Tasks Implies Structured Subspaces

Before performing numerical optimisation, we now show that—due to the heavy constraints place on the representation by eqs. (1) and (2) place—the representation, ***r***(*s, e*), *must*, in many of the cases we study, contain a set of (potentially-aligned) subspaces, each encoding a memory a particular action away. We develop an intuitive argument here, and go through the details in appendix B.1.

Begin by considering the neural subspace spanned by the read-out weights, ***W***_out_. Two things must be true about activity in this subspace, first it must encode the current observation to permit recall, and second it can’t encode the other memories as they would corrupt recall, fig. 3A. The other memories must be stored elsewhere in the representation, orthogonal (hidden) to the readout weights, ***W***_out_.

**Figure 3.**
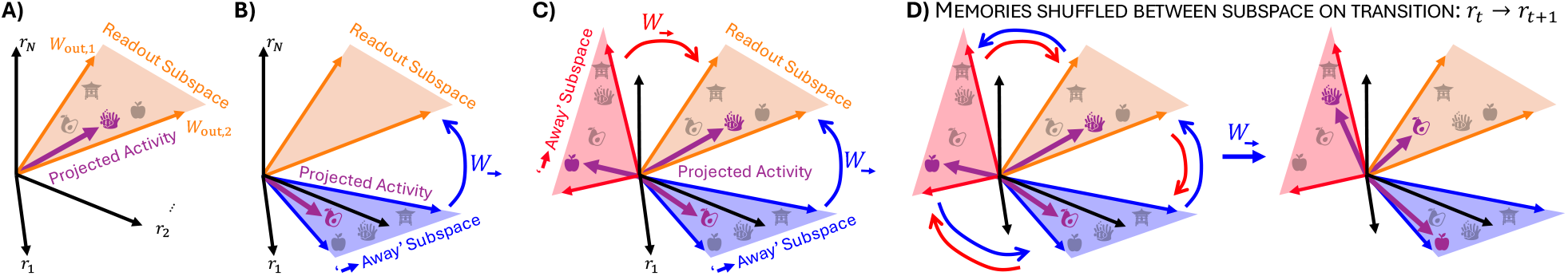
Structured subspaces are required. Examples shown for the task in fig 2D. **A)** In high-dimensional neural space, projecting the neural activity into the readout subspace (spanned by the rows of the readout matrix, ***W***_out,*i*_) must recover an encoding of the current stimulus, in this case, coral. **B)** Applying first 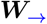 then ***W***_out_ must recover an encoding of the stimuli 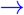-away, therefore the subspace spanned by the rows of 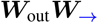, the ‘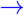 -away subspace’, must contain an encoding of the relevant stimulus, pear. **C)** Similarly, there is a corresponding subspace for the other available action, 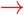. **D)** Following this logic for all possible action combinations leads to a simple representational structure: connected subspaces that each encode the stimuli at different offsets from the current position. Taking the action 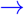 leads to a shuffling of the activity (memories) between subspaces so that each subspaces encodes the stimuli found at the subspaces’ corresponding offset from the new current position.

Now consider taking action *a* using the corresponding weight matrix, ***W***_*a*_: ***W***_*a*_***r***(*s, e*) = ***r***(*s* + *a, e*). To permit readout, eq. (2), this transformation must extract the representation’s encoding of ***o***(*s* + *a, e*) from the depths of the representation and map it to the readout subspace, so that ***W***_out_***r***(*s*+*a, e*) = ***o***(*s*+*a, e*) (see appendix B.1 for treatment with biases). Since the transformation is linear, there must exist a subspace in the hidden space where memory of the observation *a* away from your current location is stored, fig. 3B.

Applying the same logic for each possible action, there must exist subspaces encoding the stimuli displaced by each distinct action *a* (and indeed sequence of actions {*a*_*i*_} _*i*_) from your current location, fig. 3C. Each subspace answers the hypothetical ‘what would I observe if I took action *a*’. These subspaces must be interconnected by recurrent weights, ***W***_*a*_, according to the task structure. For example, taking action *a* moves the contents of all subspaces to their ‘*a*-neighbouring’ subspace, fig. 3C.

The representation could contain additional activity independent of these subspaces, but it does not help solve the task, and since neural activity is costly, the most efficient solution will remove it (though enforcing nonnegativity can change this, appendix B). As such, with these subspaces, we can summarise the underlying representation as

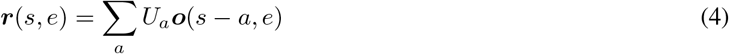

where we have defined subspace matrices *U*_*a*_ that map the observation *a*-away into neural activity. This tells us that neural activity is composed of a set of subspaces, each encoding the stimulus at a particular action-offset from your current location. The activity is then shuffled around (between subspaces) according to the rules: i.e. *U*_*a*+*a*′_ = ***W***_*a*′_ *U*_*a*_. To see that this representation, ***r***, has an associated algorithm in which memories are passed between subspaces consider taking the action *a*^′^:

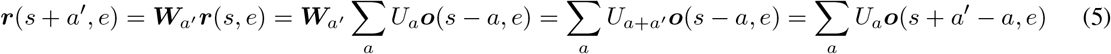

This equations tells us that, after applying ***W***_*a*_, the memories shuffle amongst subspaces according to the task rules, fig. 3D.

Finally, as long as the columns of all the subspace matrices are linearly independent, this representation solves the general structured recall task: there exists a readout matrix and a set of recurrent matrices that can correctly manipulate the activity to permit recall, independent of the particular encoded sequence. Crucially, this is true even if the subspaces are non-orthogonal (just consider a basis change to ***W***_*a*_, and correspondingly *U*_*a*_ etc., by any invertible matrix). This subspace passing algorithm is the ‘slots’ (different word, same thing) observed in nonlinear recurrent neural network models trained on similar tasks (Botvinick and Plaut; Whittington et al.; Wang, Fusi, and Stachenfeld; Piwek, Stokes, and Summerfield) and similarly matches the working memory subspaces observed in prefrontal cortex in a variety of tasks (Xie et al.; Panichello and Buschman; El-Gaby et al.). This is despite the fact that standard nonlinear RNNs (and presumably the brain) are much more expressive, and could have learnt all sorts of different algorithms.

### 3.2 An Axis of Task Diversity: Stimuli-Correlations Determine Optimal Subspace Alignment

Having established that our optimal representations are composed of subspaces, eq. (4), using numerical simulations, we now consider how minimising energy use, eq. (3) structures them; in this section we derive a relationship between the alignment of different subspaces and stimuli-correlations, and in the following section we consider subspace sizes. For a more mathematical analysis of these effects see Appendix B.2.

In our model, the optimal subspace alignments are determined by the correlations between the encoded stimuli. Intuitively, consider a neural encoding of two variables, *x* and *y*, from a dataset D of many (*x, y*) pairs:

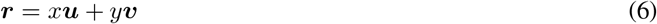

Here *x* is encoded along ***u***, and *y* along ***v***; ***u*** and ***v*** are two subspaces! Considering the firing energy of this population it becomes clear that correlated variables (memories) interact with the subspace alignment (***v***^*T*^ ***u***) to determine the cost:

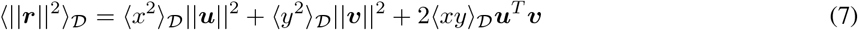

We see that the final term in the above loss depends on the correlation of the two variables (across all tasks), ⟨*xy*⟩_𝒟_, and can be reduced by anti-aligning their encodings if the variables are positively correlated, and vice versa, fig. 4B. Thus, minimising activity energy leads to negatively (positively) aligning (anti-)correlated stimuli. This is a whitening operation, a classic effect in efficient coding arguments (Atick and Redlich).

**Figure 4.**
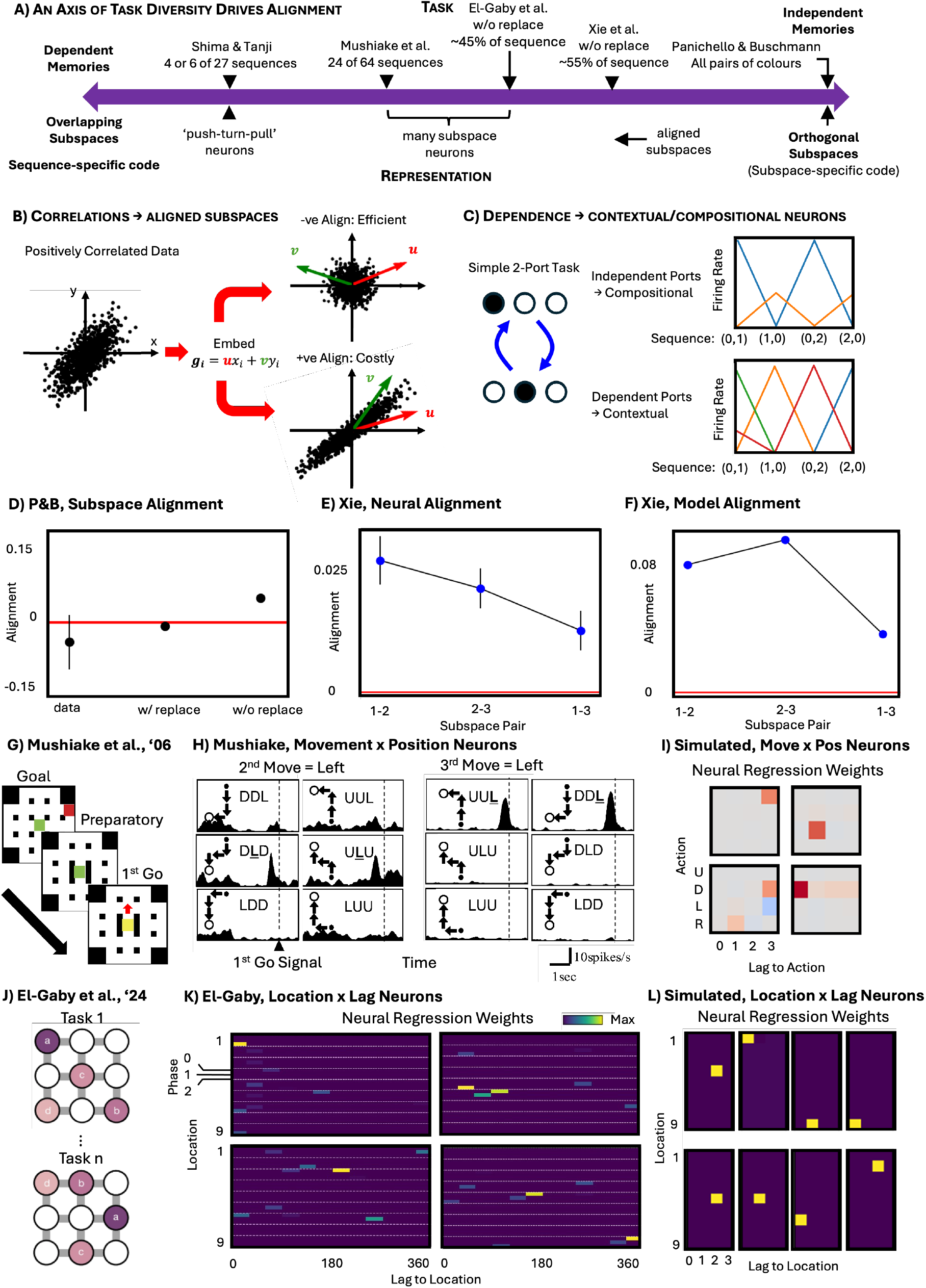
An axis of task diversity separates contextual and compositional codes and determines alignment of neural subspaces. **(A)** Optimal subspace alignment is determined by correlations in the task, sketching an ‘Axis of Task Diversity’. **(B)** Embedding positively correlated data along negatively aligned subspaces whitens the data, saving energy. **(C)** We study an agent alternating between two choices from three ports and show the neural activity of a subset of simulated optimal neurons on a subset of trials corresponding to the (0, 1) and (0, 2) port sequences. If, across all tasks the representation is optimised over, the ports are sampled independently, single neurons are compositional, e.g. the blue neuron fires for ‘Port 0 now’, while the orange fires for ‘Port 0 next’. Conversely, the optimal representation when trained on only the four sequences plotted contains mainly contextual single neurons that fire for only a single task configuration. **(D)** The subspaces in Panichello and Buschman‘s data, and our simulations of their task, are orthogonal, but sampling the colours without replacement leads to alignment, a prediction for future experiments. **(E)** Encoding subspaces in Xie et al. are aligned, **(F)** something we match in simulations. We attribute the theory-data mismatch in alignment magnitudes to estimation difficulties, see Appendix E. **(G)** Mushiake et al. train monkeys to prepare then perform sequences of three actions; they start from the centre (green square), are shown a goal (red square) and a barrier (black lines) such that only one action sequence succeeds (in this case *up, right, right*). **(H)** They report compositional neurons with delay period activity tuned to a particular action in a particular number of steps. **(I)** Top row: We find the same in simulations, though due to the correlations in the task we also, bottom row, find mixed tuned neurons. To study neural tuning we perform regression between the delay period activity and the presence of a particular action at a particular lag, and plot the weights, Appendix E. **(J)** El-Gaby et al. train mice to visit a sequence of 4 reward ports out of a set of 9. **(K)** They find compositional neurons tuned to a particular goal at a particular lag, **(L)** we find the same (plot logic as in **(I)**).

This logic means that all tasks can be placed on a continuum—an ‘Axis of Dependence’ governed by the correlations between stimuli. At one extreme, with maximal task diversity (all possible sequences), independent components are represented in orthogonal subspaces, and, if subspaces align to specific neurons, then single neurons will tuned to the the presence of a particular stimuli at a particular offset regardless of the sequence, fig. 4A. Indeed, under a slightly simplified version of the loss, we prove that this will be the case, appendix B.3. This matches the measured ‘compositional’ coding strategies, that use orthogonal or near-orthogonal subspaces for each stimuli (El-Gaby et al.; Xie et al.; Panichello and Buschman; Mushiake et al.; Basu et al.).

However, the more correlated the stimuli are (i.e., when not all possible sequences are presented leading to reduced task diversity), the more their encoding subspaces optimally align. In extremis, stimuli are dependent—knowing *x* determines *y*. As such, an encoding of *x* in the readout subspace is sufficient for the dynamics to predict *y*, removing the need for activity outside the readout subspace. This produces a purely contextual code in which single neurons code for your current state and environment, fig. 12. Tasks towards this end of the spectrum, with highly correlated stimuli have optimal representations in which encoding subspaces are highly aligned, and the single neuron responses are similarly contextual. In particular, they feature splitter-cell-like neurons that fire to a particular sequence, fig. 4C. This matches the contextual codes measured in some structured working memory tasks, such as Shima and Tanji who measured neural representations of action sequences like ‘push-turn-pull’. In our framework, Shima and Tanji measured contextual neurons because of the low task diversity, which introduces large correlations between sequence elements. In this task, knowing the first sequence element is ‘pull’ restricts the animal to one of two sequences. Had the animal experienced all 27 possible sequences, we predict the optimal representation would encode each sequence element in an orthogonal subspace.

This axis of task diversity can be used to explain a further puzzling discrepancy in subspace alignments. Panichello and Buschman present two colours to a monkey, who later has to recall one of them, and find orthogonal subspaces encoding each colour during the delay period before the animal knows which colour to recall, fig. 4D. Xie et al. present three spatial cues to a monkey, who then has to recall them in order, and find slightly-aligned subspaces that encode each stimuli during the delay period^3^, fig. 4E. Why are the subspaces orthogonal in Panichello and Buschman but slightly aligned in Xie et al.? In our framework this arises from the different task statistics. Panichello and Buschman sample colours independently (i.e., maximal task diversity), so they should optimally be encoded in orthogonal sub-spaces, fig. 4D. Conversely, Xie et al. sample stimuli without replacement (i.e., reduced task diversity), introducing correlations between sequence elements, thus aligning the optimal representations, fig. 4F.

Moreover, the theory doesn’t just predict the non-orthogonality of encoding subspaces in Xie et al. Subspaces can be aligned or antialigned, depending on whether the encoding of the same stimulus in different subspaces point in the same or opposite directions, and our theory clearly predicts which form of non-orthogonality the data should show. I.e. sampling without replacement introduces a negative correlation between memories, so we predict, and confirm in data, that the encoding subspaces are slightly *positively* aligned, eq. (7), fig. 4E-F. These arguments make testable predictions for both tasks: switching between sampling with or without replacement should swap the memory subspace orientations between orthogonal and aligned, and, in general, the structure of alignments should be predictable from stimulus correlations.

Finally, to understand the single neuron correlates of these effects, we constrain the neural activity to be nonnegative. This complicates the mathematical analysis of the optimal representations. In a simpler related setting, our previous work found that independent memories should be encoded in different neurons (Whittington et al.; Dorrell et al.; Dorrell, Latham, and Whittington). In this work, using a slight simplification of the loss, we find the same is true: subspaces encoding independent memories are optimally represented by different neurons B.3. This bolsters our numerical results regarding the single-neuron correlates of subspace alignment: aligned subspaces manifest as contextual neurons, while orthogonal subspaces are implemented by neurons tuned to a particular subspace, fig. 4C. This matches some reported single neuron tuning curves; for example, as discussed, Shima and Tanji report single neurons tuned to a particular sequence rather than a subspace. Conversely, Mushiake et al. trained monkeys to plan sequences of three actions through a grid world, fig. 4G, and observed, during a delay period, individual dlPFC neurons that code for the first action, others the second, and some the third, fig. 4H. Similarly, El-Gaby et al. trained mice to visit a sequence of four locations, fig. 4J, and found that some neurons were tuned to the location the mouse would visit next, others to the one after that etc., fig. 4K. In our theory, the degree of mixed-selectivity roughly correlates with the degree of alignment (Dorrell et al.). As such, relatively independent memories as in El-Gaby et al. and Mushiake et al., will lead to a propensity for many modular neurons, fig. 4I,L. It also interesting that not all neurons reported in El-Gaby et al. are modular, suggesting it indeed lies somewhere on this axis away from the fully modular end.

However, when additionally constraining representations to be nonnegative, our theory does not uniformly match data. Panichello and Buschman report orthogonal subspaces containing independent memories, which in our theory would manifest in disjoint single neurons. Yet the data shows no significant single neuron tuning to particular memories. It is clear from this, and other signals, section 5, that a complete theory of single neuron responses in prefrontal working memory representations remains at large.

## 4 Distance from Recall Determines Subspace Sizes and Permits Algorithmic Inference

The next natural question regarding a subspace-based solution concerns the sizes of the encoding within each subspace, i.e. the average amount of neural firing used by the encoding in each subspace. This similarly displays some puzzling patterns. Panichello and Buschman find that the two colours are encoded with equal activity, while Xie et al. find different sizes (first stimuli encoding *>* second *>* third) — a discrepancy we again confirm from reanalysis of the data, appendix E. In this section we outline how our normative framework explains this discrepancy, and use these findings to infer the presence of a ‘conveyor belt’ algorithm in the data of Xie et al. and Chen et al.

### 4.1 Optimal Sizes Follow Recall Distance

In our model the three energy losses, eq. (3), have different effects on subspace size (again, for a slightly more mathematical version of these trends, see appendix B.2):

- Activity Loss: By seeking to shrink neural activity, encourages all subspaces to be small.
- Readout Weight Loss: The smaller the activity in the readout subspace, the correspondingly larger the readout weights have to be to compensate, and vice versa. Therefore, the readout loss encourages the readout subspace to be large.
- Recurrent Weight Losses: The recurrent weights shuffle neural activity between the subspaces. If the sub-spaces have different sizes this requires the recurrent weights to grow and shrink the encodings as they move around. This is more costly than keeping the encodings the same size, fig. 5A, and so the recurrent weights exert a force towards keeping the subspaces the same size.

**Figure 5.**
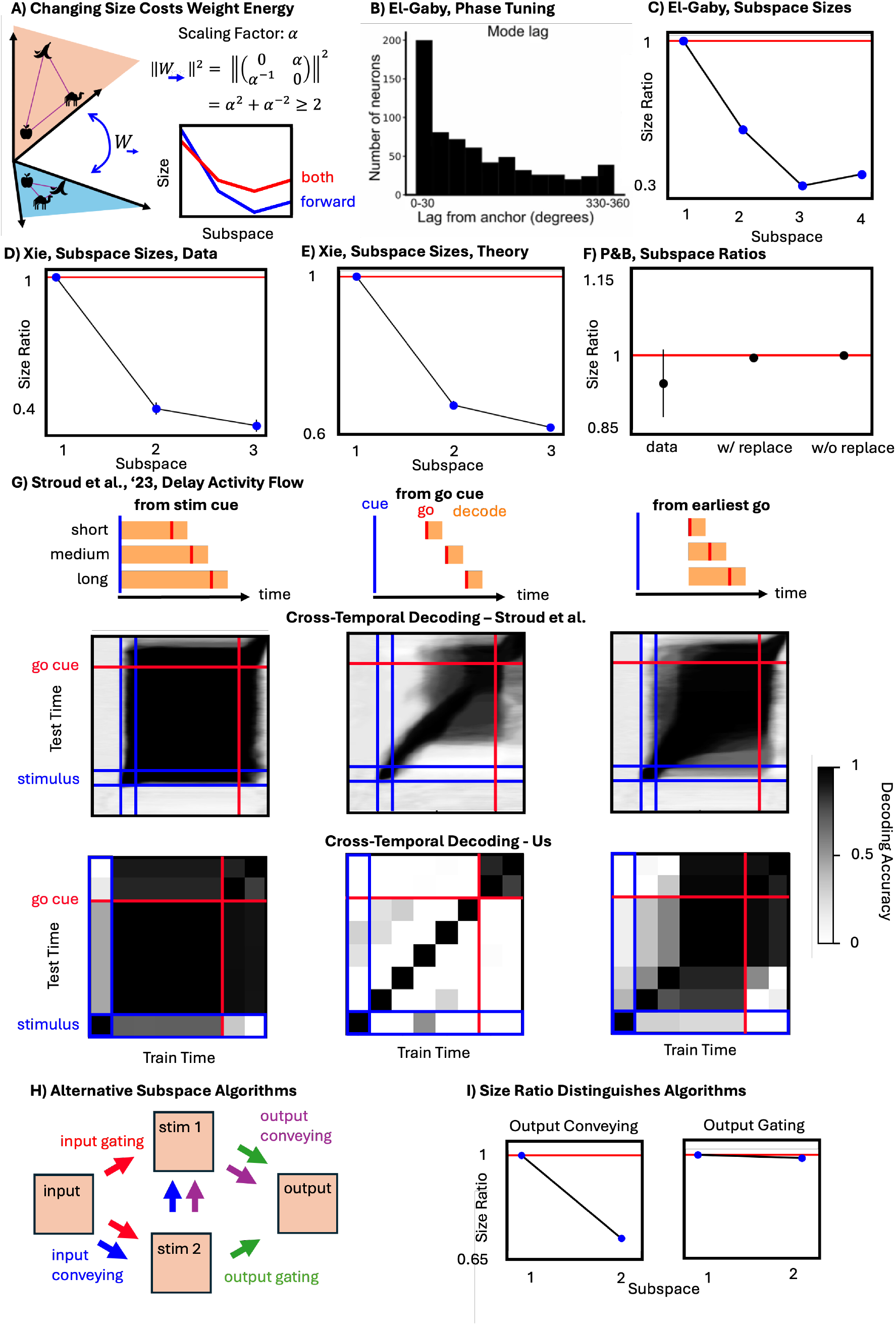
Subspace sizes are determined by their ‘closeness’ to the readout subspace. **(A)** Changing the encoding sizes between subspaces incurs a recurrent weight cost; for example, even in the simplest case where 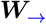 simply scales by *α*. The effect is also action dependent, a simple four memory task, as in El-Gaby et al., shows either symmetric or asymmetric size curves depending on whether both *forward* and *backward* (symmetric) or only *forward* (asymmetric) actions are included. **(B)** El-Gaby et al. report a decay then rise in number of neurons encoding memories at each lag, **(C)** we find the same in simulations. **(D)** In Xie et al.‘s data the later memories are more weakly encoded, **(E)** we find the same in theory. **(F)** Panichello and Buschman‘s data has two equally sized subspaces, an effect we recover independent of simulated sampling scheme. **(G)** Stroud et al. consider a working memory task with variable delay period and different recall requirements: the memory must either be recalled from presentation until go cue, from the go cue, or from the earliest potential arrival of the go cue. They find the representation rotates until recall is required, when it stabilises, as shown by cross-temporal decoding. We find the same in our representations. **(H)** In Chen et al.‘s data, stimuli-encodings arrive and are gated into the correct subspace (red arrows) depending on whether they arrived first or second, rather than following a fixed ‘conveyor belt’ (blue arrows). Two different output subspace algorithms are then possible: either the memories remain in their subspace until recall - output gating (green arrows), or they follow an output conveyor belt (purple arrows) implying that the memory of the second stimulus briefly appears in the first subspace. **(I)** In our theory, only the output conveyor belt solution is consistent with the observed difference in encoding sizes (Xie et al.; Chen et al.).

These effects compete; in all cases the readout subspace will be the largest, while the other subspaces will settle on a compromise solution depending on the task structure. To see some examples, consider the looping *ABCD* task with a single *forward* action. It is worth paying some recurrent weight cost to shrink the neural activity, and then slowly grow it back to full size over two steps leading to a graceful decay curve, with the memory encodings getting smaller further from readout, fig. 5A. Here all the subspaces have different sizes (fig. 5A, ‘forward’). These effects, however are highly dependent on the structure of the task. If we now consider a task with two actions, both *forward* that loops *A* →*B* →*C* →*D* →*A* and *backward A* →*D* →*C* →*B* →*A*, then subspace sizes still fall with distance from the readout subspace, but now they are symmetric (fig. 5A; the 2nd and 4th subspaces now have equal size as they are both one step away from the readout subspace).

These kinds of effects can explain the observed patterns in the data. For Panichello and Buschman it says both subspaces should have equal subspaces since they are both one-step away from being readout, whereas for Xie et al. the subspaces should gradually reduce in activity the further away from being readout, fig. 5D-F. We see similar effects in El-Gaby et al. where subspaces encoding memories further from being ‘readout’ seemingly have fewer neurons coding for them (a proxy for the size of the subspace), fig. 5B. In this task mice run in looping patterns of 4 goal locations, with PFC neurons tuned to the future locations of the mouse. The number of neurons tuned at each offset follows a graceful decline the further in the future the position is, before rising slightly returning to the present, fig. 5B. Our simulations show a similar effect, including the slight rise in encoding strength for the most distance memories, fig. 5C. This can be understood, subspace activity gradually reduces further away from the readout (Lag 0; Subspace 1) to minimise firing energy but as the subspaces get closer again to the readout (e.g., Lag 330; Subspace 4) they increase in activity again gradually to avoid particularly large recurrent weights.

### 4.2 Rotational dynamics during delay periods are energy optimal

We have considered structured working memory tasks, but similar pressures are at play in working memory paradigms without sequential structure. Consider a task in which an animal has to produce a cued action after a fixed delay. While it might seem simpler to use a stationary memory encoding during the delay period, corresponding to a single subspace, energy can actually be saved by rotating the memory through a series of subspaces. During the early delay period the memory can be encoded with small firing rates in a correspondingly small subspace, saving energy, before being rotated into a larger subspace for readout. This scheme saves firing energy at the expense of a weight cost required to implement the dynamics and additional neurons.

Indeed this is exactly what is observed in both our simulations and monkey prefrontal cortex (Stroud et al.). A critical observation in Stroud et al. is the role of variable or fixed delay periods. A fixed delay period guarantees that the memory won’t have to be recalled for a period, during which the memory can happily rotate amongst lower energy encodings. On the other hand, if the delay period varies between 1 and 2 seconds the memory can freely rotate for the first second, but must henceforth be ready to readout, even at large energy costs. They demonstrate a theory of ‘optimal information loading’ that displays three different behaviours depending on the decoding requirements. If the network has to be able to recall the stimulus at all moments during the delay period the memory encoding is fixed, fig. 5G left, whereas if the stimulus is only required after a go cue arrives then it can freely rotate until summoned by the go cue, fig. 5G middle. However, if the memory is waiting ready to be recalled, from the earliest possible recall time it displays mixed dynamics: it begins by rotating before arriving at its final representation in time for the earliest potential recall. We find the same effects in our model, fig. 5G bottom row, unifying optimal information loading with the kinds of efficient computing we have been considering.

### 4.3 Inference over Different Subspace Algorithms

We now use optimal subspace size effects to infer which ‘subspace algorithm’ is at work in the sequence memory tasks of Xie et al. and Chen et al. In Chen et al. (up to four) items are presented to a monkey in order and then must be recalled in sequence^4^. However, the stimuli could flow through the subspaces in a few different ways, i.e., different plausible subspace algorithms. There are two types of algorithm, ‘conveyor belt’ and ‘gating’, for how the stimuli are initially inputted into the subspaces and how they are recalled (outputted).

We first consider the encoding phase. The ‘simplest’ model would be a *conveyor belt*, in which each stimulus-encoding flows through a consistent sequence of subspaces, pauses during the variable delay period, then continues after the go cue, fig. 5H blue arrows. When we model this task with a *single forward* action this is the optimal learned algorithm. However, Chen et al. find that the monkey instead uses an ‘input gating’ strategy in which each observation moves from an input subspace—a holding pen—*directly* to its respective subspace depending on whether it was the first, second, or third presented observation, fig. 5H - red arrows. If we model the task such that there are different action matrices for each time step, appendix D—as could be implemented biologically by a series of neural gates, appendix A—we recover this solution as optimal even though the conveyor option could still have been learned, fig. 13.

While it is known that PFC performs input gating rather than a conveyor input, it is unknown what happens at output. Indeed, the same choice between ‘conveyor belt’ and ‘gating’ arises for the output: do memories stay in their memory subspace until they are recalled (gating, fig. 5H - green arrows), or do they flow through one another towards the output (conveyor belt, fig. 5H - purple arrows)? Again, in our model using a single *forward* action produces a ‘conveyor belt’, while flexibly swapping action matrices produces ‘output gating’, fig. 13. Unfortunately, these options are not easily distinguished from neural data because the animal’s response is too fast to be captured by calcium dynamics. However, fortunately, with our normative theory we can see measurably different predictions from these two algorithms. Following our arguments presented earlier, subspace size is determined by ‘adjacency’ to the readout subspace and since we know that the subspaces in Xie et al. have different sizes, this implies that they have different distances to the readout subspace which is only true for the output conveyor belt subspace structure but not the ‘output gating’ structure, fig. 5I.

These results raise a question: given the fully conveyor belt model is ‘simpler’, only requiring one matrix, why does the monkey use multiple input-to-storage subspaces transformation (and potentially uses a conveyor for the output)? We suggest two reasons. First, the monkeys were trained on sequences with a variable number of stimuli, between 1 and 4. This is clunky with a full conveyor belt model as even when the sequence only has one memory it has to be conveyed all the way round to be readout. By sending the memory straight to its own subspace, ‘input gating’ avoids this. However, once the correct set of subspaces corresponding to the length of the sequence have been filled, the output conveyor is ‘simplest’, using least action matrices to convey the waiting memories to readout. A potential prediction is that if monkeys are trained on a fixed sequence length, then they learn both an input and output conveyor belt. A second reason why monkey learn ‘input gating’ and ‘output conveyor’ is that the monkeys perform the output much faster than the input is provided. Thus a simple model that requires no gating (output conveyor) might be preferred at high speeds. This could be tested by forcing the monkey to perform actions at a particular, slower, frequency.

The recent results of Chen et al., along with Xie et al., Panichello and Buschman, and El-Gaby et al., show that different subspace algorithms are taking place in PFC. In this section we have shown that our general efficient computing principles can be used to reason about each algorithm, and how it should manifest in neural activity—allowing us to infer algorithm from otherwise ambiguous data.

## 5 Discussion

Here, we developed an efficient computing approach to normatively reason about prefrontal structured working memory representations. We used this approach to sketch an axis of task diversity, that unifies contextual and compositional codes as two extremes of a spectrum of optimal solutions, section 3.2. Further, we were able to normatively explain intricate measured discrepancies via subtle task differences section 3.2 and section 4.1. Finally, we used normative reasoning to predict which of two subspace algorithms underlie current recordings section 4.3. Hence, this theory allowed us to reinterpret existing measurements, understand precise representational properties, and make novel computational predictions within a unified normative framework.

This representational logic does not just apply to prefrontal structured working memory recordings; rather, we expect it to be useful in similar subspace computations. For example, the motor cortical literature often decompose activity into preparatory and potent subspaces (Churchland and Shenoy). In fact, the literature has recently described a puzzle: preparatory and potent subspaces are observed to be orthogonal, yet the same is not true of RNNs trained on delayed-reach tasks similar to those used with monkeys (Elsayed et al.). Yet, subsequently, Zimnik and Churchland found that training an RNN to perform two reaches in sequence, one after the other, led it to orthogonalise its preparatory and potent subspaces; why? This makes sense from our theory. If the network only ever performs single-reach tasks the preparatory and potent subspaces are never jointly occupied, the activity in the two subspaces is therefore highly correlated, and they align. Alternatively, dual reach tasks ensure the preparatory subspace must contain an encoding of the second reach while the potent subspace is performing the reach. If the two reaches are independent, the two subspaces should orthogonalise, as observed. Thus, we hope that our theory will be useful wherever neural subspace algorithms are a useful way of thinking about the representation.

More broadly, these subspace algorithms are intriguing. By manipulating subspaces independent of their contents they neurally implement a function. We suspect this flexibility will make them an important mode of neural computation, and hope that our normative theory will be useful in reasoning about their implementations in both brains and RNNs.

That said, our framework is clearly limited, even within the scope of working memory problems. El-Gaby et al. demonstrate that their structured working memory representation lives on top of an encoding of task phase—every neuron is tuned to percentage progress towards goal, on top of which we observe the reported future-goal encoding. We are not currently able to normatively explain this. Similarly, our model predicts that orthogonal subspaces, as in Panichello and Buschman, should be encoded in different neurons, but they are not. These are smoking guns which future work could usefully attempt to match through refinements to the theory.

More expansively, our framework is poorly placed to answer many naturally arising questions. What determines which subspace algorithm the brain uses when there are multiple available, section 4.3? How are the shufflings of these subspaces combined with meaningful computations within the subspace, such as sensory-motor transformations (Stokes et al.)? What determines when to ‘chunk’ the memory codes into hierarchical segments (Soni and Frank)?

Despite this list of shortcomings, our simple theory can explain otherwise puzzling parts of neural data. Further, the basic framework, an efficient computing hypothesis, is identical to a theory of path-integrating representations, such as grid cells (Dorrell et al.). We hope that ‘efficient computing’ approaches, by framing interesting computations in mathematical tractable problems, will be useful in reasoning about the implementations of arbitrary algorithmic processes, both hierarchical and recursive, helping us to better pull computational meaning from neural recordings.

## Code

https://github.com/WilburDoz/Prefrontal_Structured_Working_Memory_Representations

## Acknowledgements

We thank Kris Jensen & Nish Patel for reading earlier drafts of this paper. We thank Surya Ganguli for advice on subspaces and Jakub Wornbard for very generous help with subspace estimation techniques. We are very grateful to the authors of Panichello and Buschman and Xie et al. for freely sharing their data with us, without which this science would have been significantly harder.

We thank the following funding sources: Gatsby Charitable Foundation (GAT3755; W.D., P.E.L, & T.E.J.B); a Wellcome Principal Research Fellowship (219525/Z/19/Z; T.E.J.B); Gatsby Initiative for Brain Development and Psychiatry (T.E.J.B); the Jean Francois and Marie-Laure de Clermont Tonerre Foundation (T.E.J.B); Sainsbury Wellcome Centre’s core provided by Wellcome (219627/Z/19/Z; T.E.J.B); Sir Henry Wellcome Post-doctoral Fellowship (222817/Z/21/Z; J.C.R.W); European Research Council Starting Grant (NARFB/101222868; J.C.R.W).

## A Biologically-Plausible Gated Recurrent Linear Network Reformulation

In this appendix we link our theoretical framework, section 2, to gated linear recurrent neural networks. In so doing, we demonstrate a biologically plausible implementation of action-dependent recurrent weight matrices which matches simple recurrent network models of thalamo-cortical loops (Logiaco, Abbott, and Escola) and the fly ring attractor (Zhang, Wu, and Wu).

### Computing Constraint is equivalent to a gated linear RNN with connectivity defined by the task structure

To see this link, imagine an RNN that has different recurrent weights, 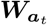, for each action. At each timestep it sees an observation, ***o***_*t*_ and if it has already seen this observation it ignores it and carries on, else the observation is new and needs to be stored. We therefore introduce a gate that determines whether this state has not been visited before, which tells the representation whether to ignore this observation or store it via a read-in matrix ***W***_0_. Mathematically:

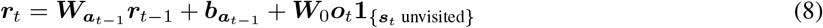

Where 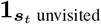 is 1 if ***s***_*t*_ has not been previously visited, and 0 otherwise. This means the RNN stores each distinct observation once. In order to do this the RNN has to have at least (number of states) × (dimensionality of observations) neurons. The RNN makes next time-step predictions via first taking an imagined step into the future and then reading out via a readout matrix ***W***_out_:

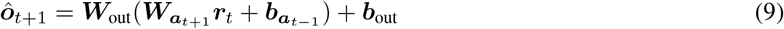

If the RNN has solved the task then, as in our theoretical representations, ***r***_*t*_ must at least be a function of the current state, *s*_*t*_, and the state-stimulus pairings given by the current environment; ***r***(*s*_*t*_, *e*). It could have more complicated dependencies, for example, it could encode each visit to the same state with different representations, but, since the task doesn’t demand it, this would be energetically wasteful.

Since this gated linear RNN uses action dependent matrices, it’s representations are constrained by the computable constraints we framed earlier, eq. (2). Thus our efficient computing problem describes the same sent of optimal representations that the above RNN would learn, but it automatically makes the activity as a function of state, *s*, as opposed to time, *t*. The benefit of posing the problem as constraints is that it lends itself to mathematical analysis, which we will use to generate our neural predictions.

### Action-Dependent Weight Matrices are Biologically Plausible

Naively, action-dependent weight matrices imply that the synapses between neurons are changing strengths based on the current action. Fortunately models have been developed, originally to study thalamo-cortical interactions (Logiaco, Abbott, and Escola), but also matched to the fly ring attractor (Zhang, Wu, and Wu), that plausibly implement such an effective synaptic connectivity change using activity. In this model a set of state encoding neurons are modelled as a linear RNN, bidirectionally and linearly connected to a second set of action-coding neurons (originally posed as the thalamus (Logiaco, Abbott, and Escola)). These action-coding neurons are not recurrently connected amongst themselves, but they are gated on or off by the current choice of action. To make this precise, imagine there were two groups of action-coding neurons, ***t***_*a*_ and ***t***_*a′*_ . Having chosen an action, one group, corresponding to the chosen action, will be free to evolve according to the linear dynamical system, while the second will be turned off. In maths, denote the state activity ***r***, the action activity ***t***_*a*_ and ***t***_*a′*_, the recurrent state connectivity ***J***, and the weights from ***r*** to each action-coding group ***A***_*a*_ and ***A***_*a′*_, and the weights from each action group back to ***r*** as ***B***_*a*_ and ***B***_*a′*_, then the linear dynamics become:

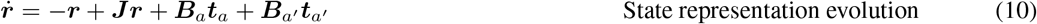

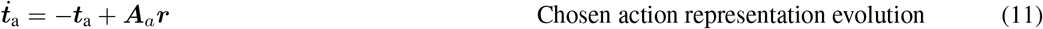

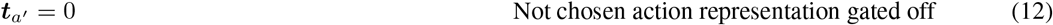

Because the action neurons are not themselves recurrently connected, their dynamics will settle to equilibrium much faster than the state dynamics. As such, we can assume ***t***_*a*_ reaches it’s steady state ***t***_*a*_ = ***A***_a_***r*** before ***r*** changes significantly. The ***r*** dynamics then become:

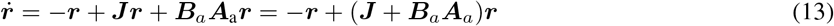

Thus there are different effective recurrent connectivities in our population through the different weight matrices (***J*** + ***B***_*a*_***A***_a_). The choice of gating can then correspond to the current action. Thus using action dependent matrices (as in our optimisation problem) has an equivalence to how actions are believed to integrated in biological neural networks.

## B Mathematical Results

In this section we derive some simple mathematical results. First, in section B.1, we show that, in certain conditions, the optimal representation has the stated subspace structure eq. (4). Then, in section B.2, we show that we can rewrite the loss as a function of only the subspace structure, and we use this to derive some simple trends. Finally, in section B.3, we introduce a simplified form of the recurrent weight loss with which we prove two things which empirically match the behaviour of the full loss: first, without nonnegativity, that independent memories are optimally encoded in orthogonal subspaces and correlations align the subspaces; second, that with nonnegativity, range-independent memories are encoded in different neurons (and hence, orthogonal subspaces). We begin with some notation.

### Some Bookkeeping

We are optimising the representation ***r*** in every state and all environments, and so let us stack all these vectors into a large matrix:

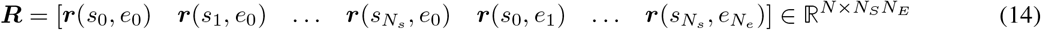

where *N*_*E*_ and *N*_*S*_ are the number of environments, and the number of states each in each environment.

We will find that, section B.1, in some settings, the representation is composed of a set of subspaces, potentially with an added bias. Using ‘matlab matrix notation’:

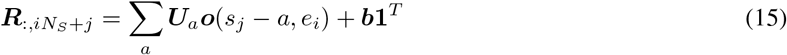

To make the notation easier let’s turn this into a single matrix vector equation with the following definitions:

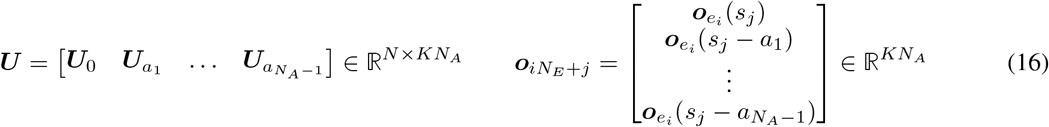

where *K* is the dimensionality of the subspace, i.e. we stack up all the subspaces and all their corresponding observations. Then simply 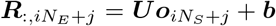. Further we can stack up all the observation vectors from the *N*_*S*_*N*_*E*_ different conditions into one matrix ***O***:

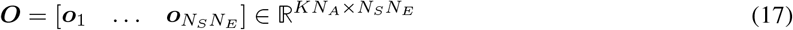

Then even more simply ***R*** = ***UO*** + ***b*1**^*T*^ . Further, we will find that a convenient variable is the subspace dot product structure: 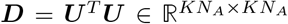. We will find we can write each of the losses in terms of this subspace dot product matrix, section B.2.

Throughout, we will assume that the encodings in each subspace are full rank and mean-zero. If this is initially not the case, enforcing it is a simple transformation. Further, we will assume that the stimuli in different positions vary independently enough that the stimuli in one subspace cannot be used to predict that in another (for example, a rule such as ‘stimuli separated by one step east are always identical’ would not be allowed).

### B.1 Subspace Structure in Representations

In this section we demonstrate that, in some settings, our optimal representation is necessarily equal to a sum of subspaces, eq. (4), matching the findings in PFC and forming the basis of our further investigations. We consider three settings:

1. A representation satisfying only the functional, eq. (1), and computing, eq. (2), constraints, and minimising the energy use loss, eq. (3). We find these representations have a subspace structure.
2. A representation in which there is an additional affine input constraint, more like an RNN.
3. A representation that additionally has both an affine read-in transformation and is nonnegative. We find that under some restrictions, this maintains the subspace structure.

#### B.1.1 No Input Constraint and No Nonnegativity

The simplest setting is one in which a representation is subject to readout, eq. (1), and computing, eq. (2), constraints, and minimises the energy loss, eq. (3), but is otherwise unconstrained. In particular, it can be any nonlinear function of the data that it wishes, and can have both positive and negative firing rates. Despite this freedom, we find all optimal representations are composed of only a set of subspaces.

First, the two constraints imply that the (mean-zero) encodings of all stimuli can be linearly decoded from the demeaned representation, 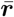:

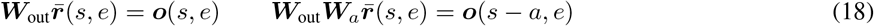

We have de-meaned the representation since the true readout and computing equations are affine, meaning they can correct for a constant shift.

This means that the full representation must be equal to the suggested subspace structure, eq. (4), plus some additional activity, ***r***_⊥_(*s, e*):

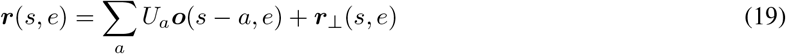

Let’s use the matrices we introduced earlier to write this as a matrix equation:

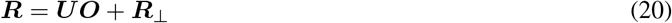

***R***_⊥_ is orthogonal in the following sense:

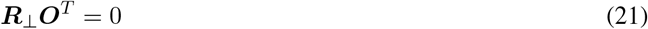

If this is not true, ***R***_⊥_ contains some encoding of the observations, which we can simply remove and shift it into our definition of ***U*** . In other words, this constraint says that the subspaces capture *all* of the encoding of the relevant observations, leaving none in ***R***_⊥_.

The second constraint is that the columns of each ***U*** and ***R***_⊥_ must be linearly independent, to ensure decoding can be performed.

Without additional constraints, this perpendicular activity plays no useful role. To make this precise, we will now examine each of the three losses and show that they are minimised by setting ***R***_⊥_ to zero. This shows that all optimal solutions can be described entirely in terms of the subspaces. Unfortunately, this approach does not generalise to the case where we enforce nonnegativity, since then ***R***_⊥_ can play a useful role in preserving nonnegativity.

##### Activity Loss

The activity loss is proportional to:

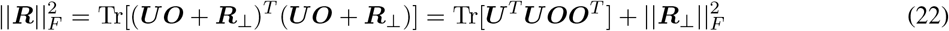

where 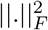 denotes the frobenius norm, the sum of the squared elements, and we have used the orthogonality of ***R***_⊥_. Hence, for a fixed ***U***, setting ***R***_⊥_ to zero is the unique minimiser of this loss.

##### Readout Loss

The min-norm ***W***_out_ matrix accesses only the correct readout subspace. We can break it into two components, one that lives within the span of the subspaces, ***U***, another that lies orthogonal to this:

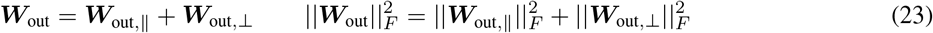

The component within the span of the subspaces is completely constrained by it’s behaviour on ***U*** :

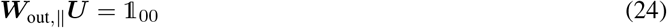

where 1_00_ is a matrix with identity in the first *K* × *K* block and zeros elsewhere.

The perpendicular component cancels the remaining impact of the demeaned 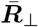:

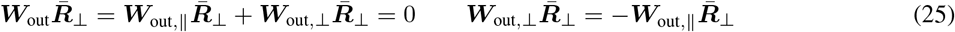

Any operation that orthogonalises ***U***_0_ and 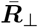. will minimise this loss for a fixed ***U*** . One example is ***R***_⊥_ = 0.

##### Recurrent Loss

This follows a very similar argument to the previous section. Each min-norm recurrent matrix can again be broken into a component within the span of the subspace matrix, and a component orthogonal:

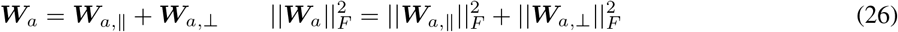

Since ***R***_⊥_ ***O***^*T*^, the perpendicular activity cannot be used to aid the help with the swapping of the subspaces. Hence, the component within the span of the subspaces must perform the relevant permutation:

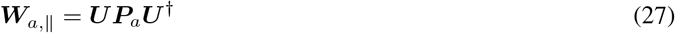

where ***P***_*a*_ is the permutation matrix that appropriately shifts subspaces to their destination under-*a* (see eq. (80) for an example).

The perpendicular component has two jobs, one is to correct for the overlap between the subspaces and 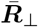:

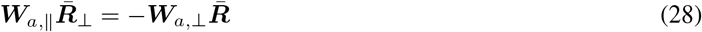

The other is to perform the correct recurrent updating of 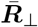. Both of these are energetically costly and not required: again, we see that we can minimise this loss by setting 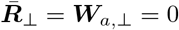.

This completes our result: the min-energy solutions will all contain only a subspace structure.

#### B.1.2 Linear Input and Transformations

In this section we consider a model where we specify a linear read-in operation, by which the memories arrive in the representation. We combine this with a linear readout, eq. (1), and computing, eq. (2), constraints to find that the neural activity is made of a series of subspaces, each encoding the stimuli at a particular offset from the agent’s current position. We will return to affine transformations in the next section.

On the first timestep this agent arrives in the world and receives an input about the current state’s stimulus. The activity after one step will be:

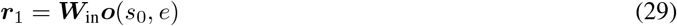

Since the set of observation vectors {***o***(*s, e*)} _*s,e*_ span a *K* dimensional space, the encoded observation lives within a *K* dimensional subspace in neural activity spanned by the columns of ***W***_in_.

After an action *a*, a second observation arrives, and the representation updates itself:

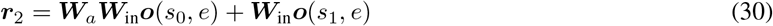

where *s*_1_ = *s*_0_ + *a*. Now the first memory has moved into a subspace spanned by the columns of ***W***_*a*_***W***_in_, while the new memory ***o***(*s*_1_, *e*) lives in the subspace spanned by the columns of ***W***_in_. To simplify notation let’s define a new set of subspace matrices, ***U***_0_ = ***W***_in_, ***U***_*a*_ = ***W***_*a*_***W***_in_, ***U***_2*a*_ = ***W***_*a*_***W***_*a*_***W***_in_ and so on.

We see that as the agent travels from state to states encoding each observation into its neural activity, ***r***, the memories are shuttled from subspace to subspace. For this to work correctly, the action matrices must obey certain constraints. For example if one action, *a*, is the inverse of another, *a*^*′*^, meaning (*s* + *a*) + *a*^*′*^ = *s*, then the corresponding action matrices must be inverses of one another: 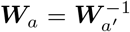, to ensure that the neural representation correctly keeps track of your state:

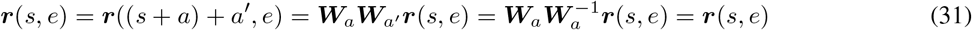

This is just the computing constraint, eq. (2), in action. In general the action matrices must match the rules of the environment. For example, if you are on a 2D grid world then moving north, then east, is the same as moving east, then north. Correspondingly, the action matrices associated with north and east must commute to ensure your representation is independent of route:

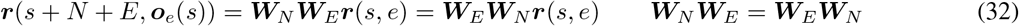

These consistency requirements on the action matrices tell us that in a working representation the action matrices will shuttle the information between the appropriate encoding subspaces.

Eventually, after all observations in the current environment have been loaded in, we have the full representation:

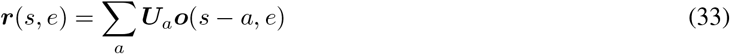

There will be one subspace for each of the relative positions (defined by some action *a* away from the current position *s*) in the environment. Note that the actions *a* in this equation are not restricted to being one step away, but can be multiple (e.g., *a*^*′*^ may correspond to 3 steps north and 1 steps east), whereas the actions in the recurrent weight matrices, ***W***_*a*_, are for one step at a time.

Once the memory representation is fully constructed it takes no further observation inputs, simply updating itself according to the action matrices which has the effect of appropriately shuffling the observations around the memory representation. For example, imagine a world with two states, and correspondingly two subspaces:

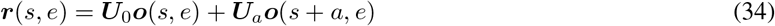

The only action available in this world is the one that swaps your state. Since swapping your state twice corresponds to staying where you are, the corresponding action matrix must square to identity: 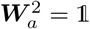. Then if you take the only action available to you the representation updates:

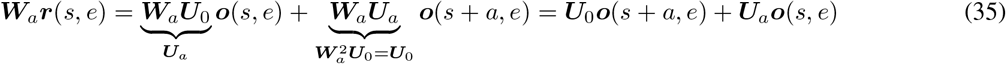

And we can see that the observations swap subspaces.

Returning to the more general case, we consider the functional role these subspaces have to play. Independent of the behaviour of all other memories we have to be able to decode the current observation using our linear readout:

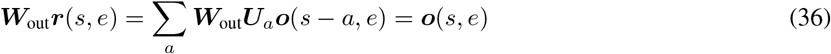

For this to be true the readout activity must perfectly extract the *K* dimensional subspace encoding the current memories: ***W***_out_***U***_0_ = 𝟙, while ignoring all the other subspaces: ***W***_out_***U***_*a*_ = **0**. This is only possible if the *K* columns of ***U***_0_ are linearly independent of the columns of each of the other subspaces ***U***_*a*_, as otherwise information contained in the other subspaces would be read-out by the read-out matrix. We can extend this argument further, applying first an action matrix and then the readout matrix lets us make the same argument about a different subspace:

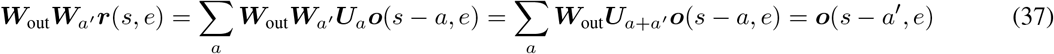

Therefore we derive a new condition on the behaviour of ***W***_out_***W***_*a′*_ ; similarly to above, it must extract the full *K* dimensional subspace encoded in subspace ***U***_−*a′*_, ***W***_out_***W***_*a′*_ ***U***_−*a′*_ = 𝟙, while ignoring all other subspaces ***W***_out_***W***_*a′*_ ***U***_*a*_ = **0**. In order to do this, as above, the columns of ***U*** _*a′*_ must be linearly independent, and linearly independent of the columns of all other subspaces.

Therefore we find, as claimed, that for the each of the *N*_*A*_ different actions, the representation needs to contain a *K* dimensional subspace, linearly independent from each of the others^5^.

#### B.1.3 Representation Decomposes even with Bias

Now we also show that the above ideas partially generalise to positive representations that require a bias, for example representations that are nonnegative. We assume that the set of all sequences of available transformations form a group and the marginal distribution of stimuli in each state are equal, as in, for example, the experiments of El-Gaby et al. In this setting we find that the representation is composed of subspaces and a bias.

Thus we return the biases to the readin and recurrent transformations, i.e.:

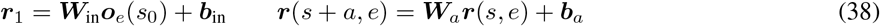

As actions are taken and new observations are seen, ***r*** now develop trailing bias terms:

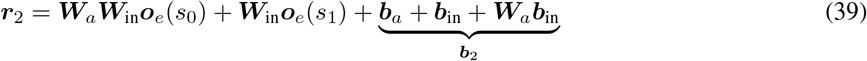

In general this bias vector will be a function of the sequence of actions taken. We develop our analysis of this trailing bias term in two separate settings: when the allowed actions do and don’t form a group.

##### Transitions around space form a group

First, we consider the case where the set of sequences of allowed transitions form a group. For example, the map might be a periodic one or two dimensional space, Figure 6a. This tells us two things: first every subspace will always contain a memory, because from every point we can move using any action and expect to find something. Second, we can legitimately keep moving in any direction for ever, by applying the same transformation repeatedly. If the length of the bias vector is not kept constant by the recurrent affine transformations then eventually, as the trajectory lengthens, it will either explode (costing infinite activity energy) or go to 0, which isn’t allowed because the representation must be positive and the only thing allowing this to be possible is the bias vector. Therefore, the length of the bias vector must be preserved under all recurrent transformations.

**Figure 6.**
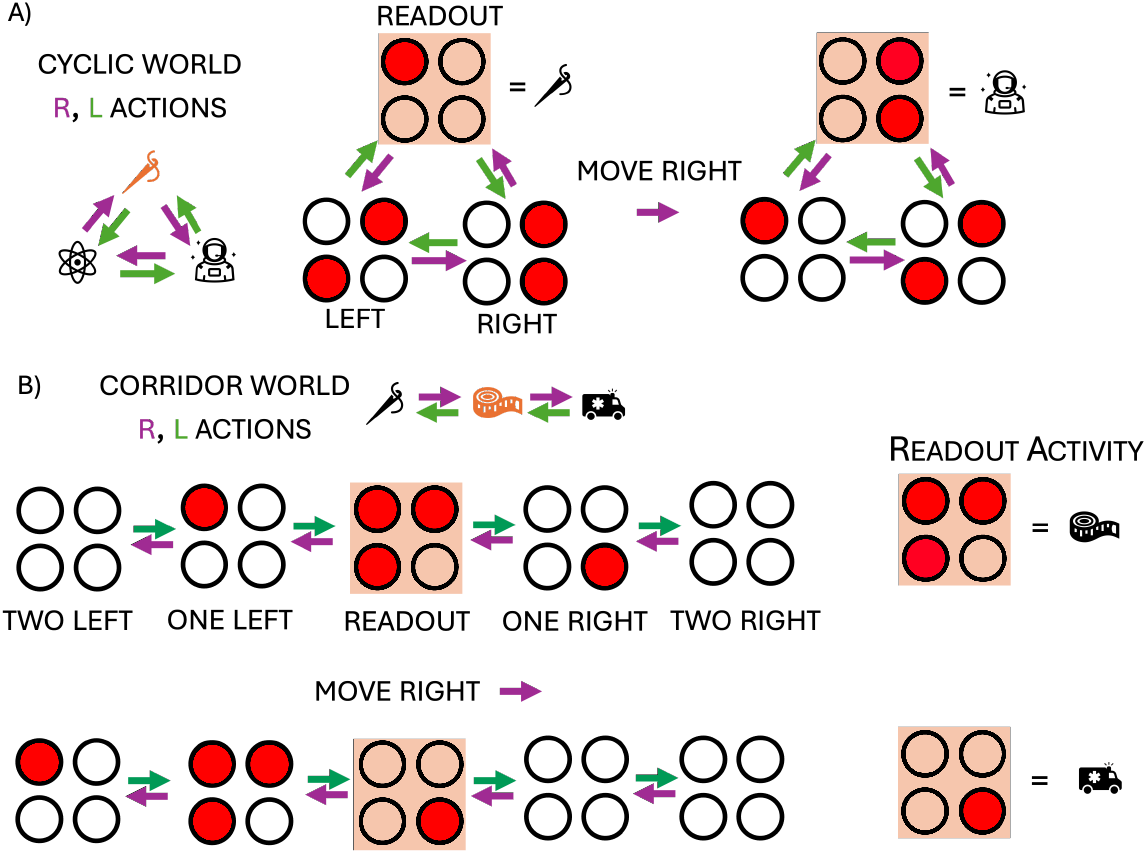
a) In a cyclic world (for example the illustrated 3-state cycle) like this you only need three subspaces, for observations at your current, left, and right positions. As you move each subspace will always contain a memory. b) However, in a corridor world you need more subspaces for the same number of states, and there will always be two empty ones.

Could the bias vector rotate though? The answer is no since, for any neuron, the bias should be as small as possible while still maintaining the nonnegativity of the representation, and rotating the bias vector increases and decreases the bias for each neuron thus costing energy. Furthermore, the bias should be action independent. This is because while there may be one environment where after taking action *a* there is less bias required in a given neuron, this will not happen across all environments since we have assumed that the marginal distributions of observations in each location are equal. Thus there will be another environment where the observations are exactly reversed, before taking action *a* a smaller bias is required, but after, a larger one. Therefore, in order to ensure the representation is nonnegative in all environments while being optimally small, the bias must be action independent (***b***_*a*_ = ***b***) and left unchanged by all affine action transformations: ***W***_*a*_***b*** + ***b***_*a*_ = ***b*** for all *a*.

Now we consider reading out from a representation, and as in the previous section we will show that the subspaces are linearly independent.

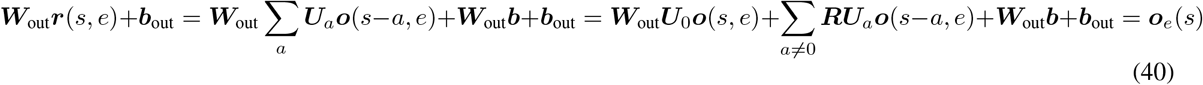

In order for this decoding to happen correctly the readout matrix must faithfully copy the coding information in the current state subspace, ***W***_out_***U***_0_ = 𝟙 and must ignore all the other information, ***W***_out_***W***_*a*_ = **0**. This is only possible if the columns of ***U***_0_ are linearly independent from all the columns of the ***U***_*a*_ matrices, as before (increasing ***b***_out_ has no cost, so it doesn’t matter whether ***W***_out_ is orthogonal to ***b***, since we can simply choose ***b***_out_ = − ***W***_out_***b***). And, as we did in the previous section, we can extend the argument by considering reading out from another subspace by first taking an action, then reading out, ***W***_out_***W***_*a*_, we derive again that the columns of all the subspaces need to be linearly independent.

##### Transitions do not form a group

Things are slightly more complex when the transitions do not form a group. For example, you might be in a one-dimensional corridor of length 3, Figure 6b. If you are in the middle subspace then the ‘current position’, ‘one left’, and ‘one right’ subspace will all have encoded memories. On the other hand, if you then move to the right, a different set of three subspaces, the ‘current position’, ‘one left’, and ‘two left’ subspaces will contain activity. Further, taking the ‘move right’ action from the furthest right point has no meaning, as that’s the end of the corridor. Thus since all states are not equal, it may be optimal to change the length of the bias vector for each state. However, it will never be optimal to make it path dependent, since the aim of the bias (making the representation minimally-positive) is invariant to path used to reach a given state.

Therefore we end up with the following representation:

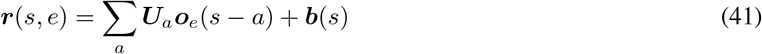

However, this does not fundamentally change the claims. In order to correctly decode the subspaces their columns must be mutually linearly independent. The only slight complication is that the representational bias, ***b***(*s*), and the readout bias, ***b***_out_, have to be chosen so that they always cancel during readout:

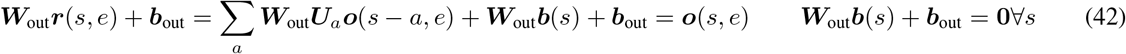

This is a minor nuisance, and so, henceforth, we restrict to group-structured settings. However we conjecture, and have empirical evidence, that our main results will generalise to cases where the transition structure is not a group.

##### Summary

In order to satisfy the constraints the representation must be made of linear subspaces encoding the observations at each relative position from the agent. If this is true there exist affine action maps (***W***_*a*_), read-in, and read-out transformations that can construct and decode the representation. We can therefore optimise the loss, eq. (3), directly over the subspace structure, potentially including the remaining positivity constraint.

### B.2 Analysis of Loss

In this section we assume the representation has a subspace structure, as found under certain conditions in section B.1, and use this to develop the loss. We find that the loss can be written entirely in terms of the subspace dot product matrix, ***D*** = ***U*** ^*T*^ ***U*** and the bias ***b***. Finally, we introduce a simplified recurrent weight loss with which we will later be able to prove some simple properties.

#### B.2.1 Analysing Losses

##### Activity Loss

The activity loss measures deviation of the firing rate from a target firing rate:

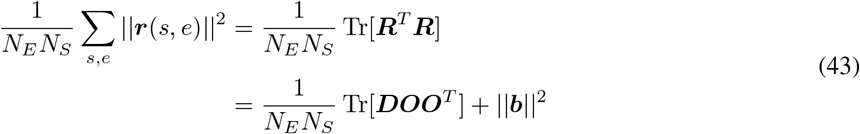

##### Readout Loss

Next, the readout matrix must simply extract the current subspace. For a given subspace structure the minimal L2 norm matrix that does this is given by the first *K* rows of the pseudoinverse matrix of ***U*** (because the first *K* rows of the pseudo inverse ‘attend’ to the first *K* columns of ***U*** which is the first subspace), denoted by 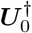, hence:\

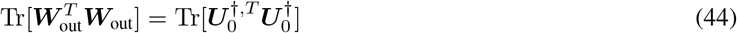

Let’s define ***D*** = ***U*** ^*T*^ ***U***, which is the subspace dot product matrix without the bias. Then, by the pseudoinverse or otherwise, one can find that ***U*** ^*†*^***U*** ^*†,T*^ = ***D***^−1^. Hence:

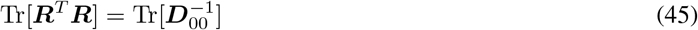

Where the subscript 00 denotes the first *K* × *K* dimensional block of the matrix ***D***^−1^.

##### Recurrent Loss

Last we have to consider the recurrent weight loss. Each recurrent weight matrix has to take the activity in one subspace and move it to its appropriate target subspace. For a given subspace structure the minimal L2 norm matrix that does this is the following:

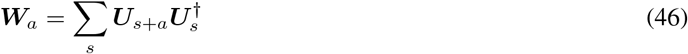

The matrix decomposes activity onto axis aligned subspaces using the psuedoinverse matrices, 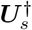, then projects them back using ***U***_*s*+*a*_. We can rewrite this using a permutation matrix, ***P***_*a*_, that maps each *K*-dimensional block *s* to the block *s* + *a*: =

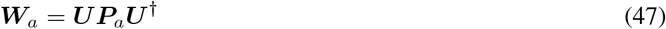

Then the weight loss is:

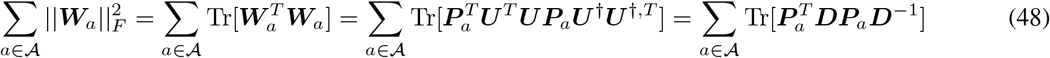

##### Combination

We can write the whole loss in terms of the subspace, and subspace-and-bias dot product matrices and problem specific parameters:

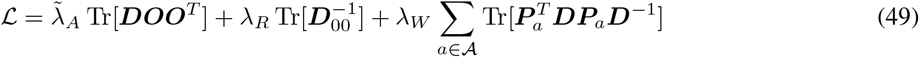

where we’ve introduced the rescaled 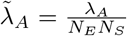. This has to be minimised over ***D*** and ***b***, subject to the representation maintaining nonnegativity and linearly independent subspaces.

##### Non-Convexity

Deriving analytic results for the optimal solution to the above equation is difficult because it is non-convex. The term that causes the problems is the recurrent weight loss. We provide code that shows that the recurrent term is a nonconvex function of the subspace similarity matrix, ***D***^6^.

#### B.2.2 Rough Trends (ignoring nonnegativity)

Without the nonnegativity constraint the bias plays no useful role, so it may be set to zero. We can then take the derivative of the loss with respect to ***D***

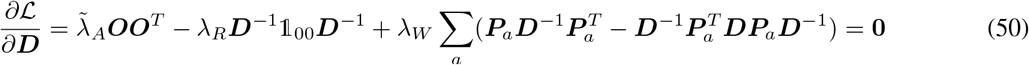

where 𝟙 _00_ is an identity matrix only in the first *K* by *K* block. From this we can derive some interesting limiting behaviour:

1. If *λ*_*W*_ is much smaller than the other hyperparameters then 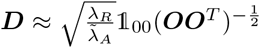. Thus the only meaningfully sized subspace is the readout one, whose dot product structure is determined by the 00th block of the observation dot product matrix. If *λ*_*R*_ is very big you want a large readout subspace so that the weight matrix can be small, while if 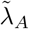 is very big you want it to be small, and to instead grow the size of the readout weight matrix. This is all intuitive: if *λ*_*W*_ is negligible then very large recurrent weights can move content between subspaces in a way that ensures there is little activity used in all subspaces but the readout one (you need large recurrent weights from adjacent subspaces to the readout subspace). This ensures that the entire subspace structure is essentially driven by the only meaningfully sized subspace, the readout subspace, whose size has to be large due to the nonnegligble *λ*_*R*_ hyperparameter. Further, if the activity in the readout subspace also became very small (due to high *λ*_*R*_) you would require very large readout weights to map the activity to the outputs.
2. If *λ*_*W*_ is much bigger than the other hyperparameters then ***D*** ≈ 𝟙 is a solution, i.e. each subspace is orthogonal and equally sized to all other subspaces. This is again intuitive: the recurrent weight matrices have to shunt activity from one subspace to another, which requires both large and small weights if the subspaces are aligned or have different sizes. Large weights cost energy. If the subspaces are orthogonal and equally sized, then the recurrent weight matrix can also be orthogonal which is its lowest energy form.
3. The subspace structure is in part determined by ***OO***^*T*^ (from the activity loss), which describes the correlations of the observations. Thus introducing correlations in ***OO***^*T*^ will introduce correlations (alignment in neural space) between the subspaces.

### B.3 Simplified Recurrent Weight Loss

The difficulty in analysing this loss comes from the (non-convex) recurrent weight loss. We therefore introduce a simplified recurrent weight loss, and use it to make two simple claims. First, without nonnegativity, independent memories are optimally stored in orthogonal subspaces, and correlations align the optimal subspaces. Second, with nonnegativity, range-independent memories are stored in different neurons.

#### B.3.1 Recurrent Weight Loss Simplification

Each recurrent weight matrix can be written as a sum of components (note this is the same as Equation (47) but where the permutation matrix is effectively performed by the sum over subspaces *s*):

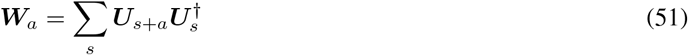

Each component extracts the activity from one subspace using 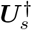, and projects it to the appropriate part of the representation with ***U***_*s*+*a*_. The Frobenius norm can then be written:

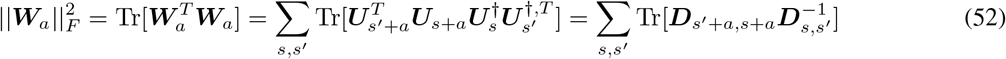

where the notation *D*_*ij*_ denotes the *ij*th *K* dimensional block of *D*. The difficulty in analysing this loss comes from the cross-terms. We therefore introduce a simplified form of the loss which does not penalise these cross terms; rather, we penalising each section of the weight matrix separately:

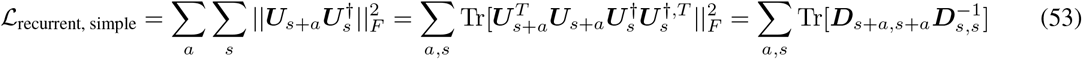

This version of the loss can be interpreted as if the movement from each subspace to another was implemented by a separate set of neurons, of the sort of ‘action neurons’ described in Appendix A. This seems quite plausible, especially if the agent needs to combinatorially combine all possible subspace-to-subspace transitions, having them independently represented might confer significant advantages. Regardless, this seems a reasonable loss to study.

Therefore we arrive at our simplified loss function:

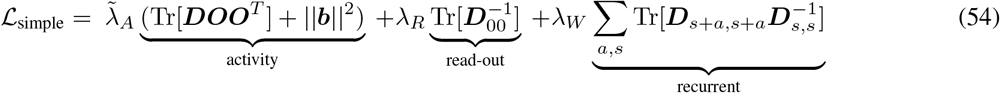

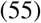

Which we can rewrite a more simply with the following notation for the block-diagonalised and shifted-block-diagonalised matrices:

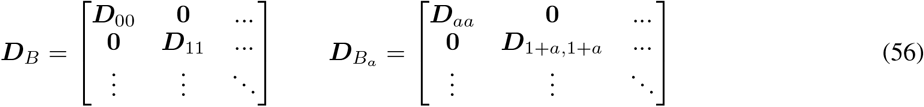

Which makes our loss:

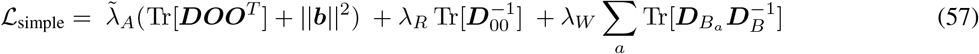

#### B.3.2 Without nonnegativity, Independent Memories are Orthogonalised, Correlated are Aligned

Here we drop nonnegativity, letting us take ***b*** = 0, and study the simplified recurrent weight loss. In particular, we start with the simplest case; when ***OO***^*T*^ takes a particularly simple block diagonal form. This occurs when the memories held in different subspaces are independent of one-another. Here we show that, in this case, the activity loss is independent of the off-diagonal blocks of ***D*** and that both weight losses are minimised by orthogonalising the slots. This means that the optimal representation for independent memories is when the subspaces are orthogonal to one-another. Conversely, non-zero off-block-diagonals in ***OO***^*T*^, i.e. correlations amongst the memories, align the optimal subspaces.

We now show that, for independent memories, each of the three losses is either unchanged or reduced by orthogonalising the subspaces. Therefore, the optimal solution uses orthogonal subspaces. Finally, we show the same is not true for correlated memories. Throughout, orthogonalisation refers to the operation of setting the off-block-diagonal components of ***D*** to zero, i.e. the within-subspace structure is preserved, but they are rotated to be orthogonal.

##### Activity Loss

It is easy to see that ***OO***^*T*^ is block-diagonal for independent memories if we consider its *ij*th K dimensional block:

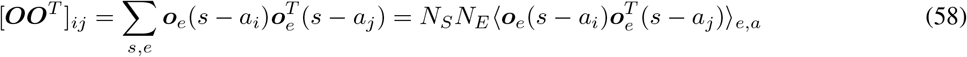

Across actions and environments the memories are independent and mean zero so this matrix is block diagonal, and the diagonal blocks are the identical covariance matrix of the observations in a single subspace:

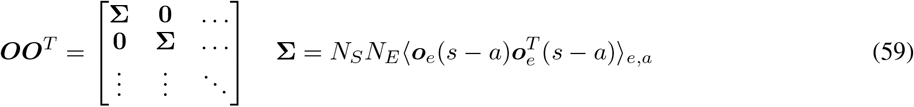

This means that the activity loss only depends on the diagonal blocks of ***D***:

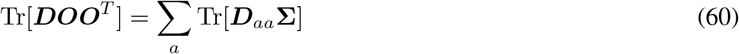

so is not affected by orthogonalisation, which only changes the off-block-diagonal parts of ***D***.

##### Readout Loss

We now consider the readout loss, and find the loss is minimised when subspaces are orthogonal. The min-norm readout matrix is equal to the first block of the pseudoinverse of the subspace matrix: 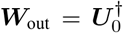. In general, 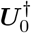 can be thought of as doing two jobs. First, within the readout subspace (the space spanned by the columns of ***U***_0_) its behaviour is to ‘undo’ the subspace embedding, so that it recovers any vector 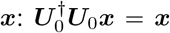

Second, it ignores embeddings in other subspaces: 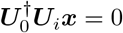 for *i* ≠0. As such, it is useful to break down 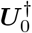 into components, one that lies within the span of the columns of ***U***_0_, and another orthogonal to all the columns:

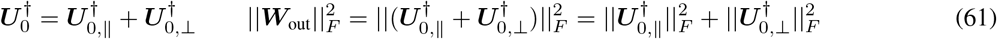

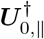 is given by the condition that:

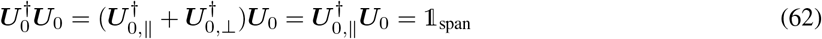

Where 𝟙 _span_ denotes identity within the span of the columns of ***U***_0_ and zero elsewhere.

Let’s consider how the readout weight loss would change if we were to orthogonalise the subspaces. This procedure leaves the within-subspace dot product matrix, 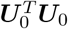, fixed. This leaves the Frobenius norm of the parallel component of the pseudoinverse unchanged, which can be seen from, among other things, the reduced SVD:

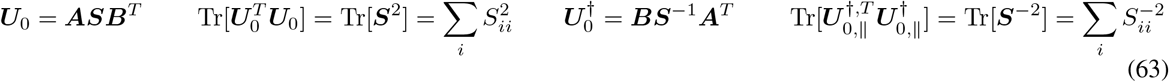

So we just need to consider the perpendicular component. This only exists to ensure that the pseudoinverse is orthogonal to all other subspaces:

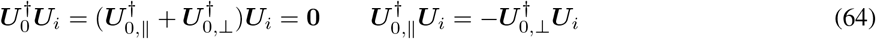

If the subspaces are made orthogonal the perpendicular component, 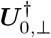, is set to zero, necessarily reducing or, in the case that the ***U***_0_ was already orthogonal to all the other subspaces, leaving invariant the readout weight loss.

##### Recurrent Loss

We now consider the recurrent loss, and observe that (due to a similar analysis to the above section) the loss is minimised when subspaces are orthogonal:

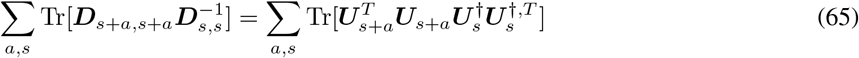

If the subspaces are orthogonal, then 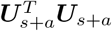 is constant. Thus we just need to consider the pseudoinverse terms. In a similar approach to the above section, 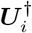, can be broken into a component within the span of its subspace, ***U***_*i*_, and one orthogonal to it, such that:

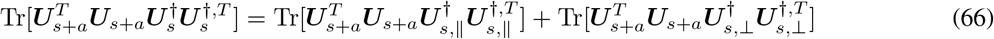

If the subspaces, ***U***_*s*_, are orthogonal, then all perpendicular parts, ***U***_*s*, ⊥_, are zero while all the parallel parts are fixed. Thus orthogonal subspaces necessarily reduces this loss.

##### Correlated Memories

Conversely, if the memories are linearly correlated across states, i.e. the matrix ***OO***^*T*^ is no longer block diagonal, we can see that the subspace dot product matrix, ***D*** won’t be block diagonal either. The activity loss will reduce if we vary the off-block-diagonal parts of ***D*** appropriately such that the off diagonal terms, such as Tr[***D***_01_(***OO***^*T*^ )_01_] are negative. Since the weight loss is minimised by orthogonal subspaces, introducing a small part of this off-block-diagonal component won’t change the weight loss too much (to first order). Therefore linearly correlated memories will align subspaces at least a bit.

#### B.3.3 With nonnegativity, Range-Independent Memories are Encoded in Different Neurons

In this section we study optimal representations for the loss with the simplified weight loss (Equation (57)), while preserving the nonnegativity constraint on neural firing. We will show that, in this setting, range independent memories— this is when all combinations of memories in subspaces are allowed but statistical dependencies may exist, i.e. the range is independent—are optimally stored in different neurons. In this setting the only appearance of the bias in the loss is in the term ||***b***||^2^, so the bias should be as small as possible, while guaranteeing the nonnegativity of the representation.

Following from a combination of our previous work, Dorrell et al., and the previous results on minimising via orthogonalisation, we consider an operation in which we break any neuron tuned to multiple memories apart. We will call this modularisation, as it corresponds to breaking the encoding of each subspace into a different ‘module’ of neurons. This operation also orthogonalises the encoding of different subspaces, therefore, as in section B.3.2, it necessarily reduces the weight losses. We show that for range-independent memories^7^ this neuron-wise modularising operation also necessarily reduces the activity loss, and therefore that the optimal representation only has neurons tuned to single memories.

Let’s say we have a representation that contains neurons tuned to multiple memories, e.g.:

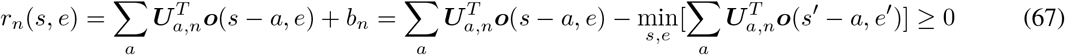

where ***U***_*a,n*_ is the *n*th row of the ***U***_*a*_ subspace matrix. Now let’s imagine breaking this into *N*_*A*_ separate, minimally nonnegative, neural tunings:

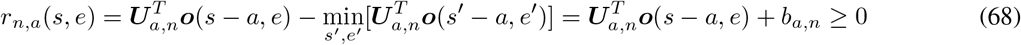

Further, we use the range independence to break apart the bias term^8^:

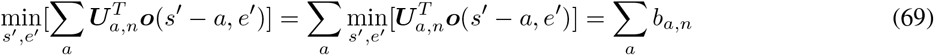

So we can actually write the neuron tuned to multiple memories as:

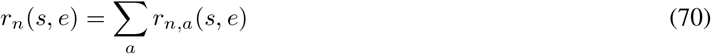

Now we can see that this operation necessarily reduces the activity loss. This is because the modular cost is ∑_*a*_ ⟨(*r*_*n,a*_(*s, e*))^2^⟩ _*s,e*_, while the mixed neural tuning can be written as the sum of the same modular cost and a postive term:

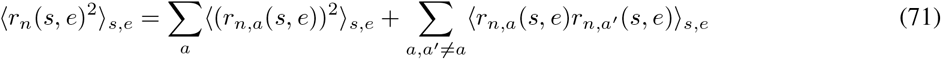

And since the variables are range independent the inequality ⟨*r*_*n,a*_(*s, e*)*r*_*n,a′*_ (*s, e*) ⟩ _*s,e*_ *>* 0 is strictly true. Since all losses are either reduced or remain constant in doing this modularising procedure, for range-independent variables there can never be a neuron tuned to multiple memories in the optimal solution.

## C L2 Regularisation as Limit of Noisy Accuracy

Here, we show that L2 regularisation on weights and activity can be derived as the limit of an accuracy loss under Gaussian noise.

Consider a neural representation matrix, ***R*** ∈ ℝ^*N×T*^, where *N* is the number of neurons and *T* the number of trials, which we linearly map through a weight matrix, ***W*** ∈ ℝ^*D×N*^, to an observation matrix, ***O*** ∈ ℝ^*D×T*^ . We consider jointly optimising ***R*** and ***W*** to linearly reconstruct the observations in the presence of additive iid zero-mean Gaussian noise on both the representation, *η*_*R*_, and weights, *η*_*W*_, with variances 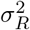 and 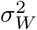.

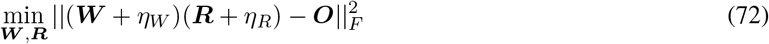

Using the properties of the noise we find the objective is:

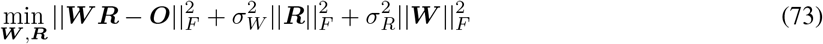

In the limit of loss noise variance, the optimisation will effectively perfectly minimise the mean reconstruction error, 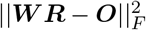, before then turning to the remaining turns, in effect giving us the following problem:

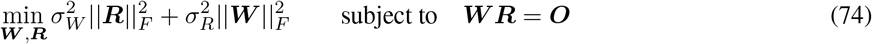

This is exactly the situation we consider: perfectly fitting data with regularisors on the L2 norms of ***W*** and ***R***. Instead, however, these are justified purely from noise and a reconstruction error.

## D Numerical Simulations

In this appendix we discuss the numerical results presented. We begin by outlining our optimisation schemes, then the different tasks. Finally, we describe the analyses applied to the optimal representations.

### D.1 Optimisation Schemes

Our simulations broadly fall into three categories. The first aligns more closely with the mathematical framing. We use the derived subspace structure to write all representations as an affine function of a set of lagged observations. We then optimise over this choice of affine function, effectively optimising directly over the ***U*** matrix. The second is similar, except we skip the affine map parameterisation, and directly optimise over the neural representation matrix itself. The third set is a gated-linear RNN: a linear RNN with action-dependent recurrent weight matrices. In this case we optimise over the weights and biases, as in a traditional RNN setup.

We largely optimise directly over the representation, since it uses fewer assumptions. However, when there is an assumed linear input or no nonnegativity, then, as discussed in section B.1, the representation has a simple subspace structure that can be used to make the simulations much more efficient.

#### D.1.1 Optimising over Affine Readin Matrix

We saw in section B.1 that some representations, after experiencing every observation, can be written as a sum of a set of subspaces, each encoding the memory at a particular offset, and a bias. In this optimisation scheme we make use of this and optimise directly over the subspace matrices.

Our full loss used to train the representation has five terms: three as in the energy loss, eq. (3), and two that enforce perfect readout, eq. (1), and path-integration, eq. (2):

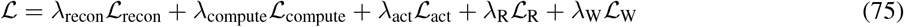

Finally, if we are including nonnegativity we additionally shift the representation up by its minimal value to make it nonnegative. We choose the values of *λ*_recon_ and *λ*_compute_ to be large enough to enforce the constraints to a high level of satisfaction. The other *λ* values are part of the problem statement, we usually set them to 1.

##### Creating the Representation

For each environment and each state we create a matrix containing the observation at each relative action offset: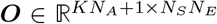, where *N*_*E*_ is the number of environments, *N*_*S*_ is the number of states, *N*_*A*_ is the number of subspaces, and *K* is the dimensionality of each subspace. The final row of ***O*** is all 1, to encode a bias. Then we parameterise the representation using the subspace matrix, 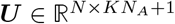 which maps the inputs into a neural representation:

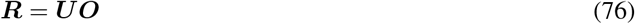

We now describe each loss in turn.

##### Readout Loss and Functional Constraint

Given a representation we construct the minimal L2 norm readout matrix. Since the readout includes a bias that is not penalised, it makes use of it to perform all parts of the readout mapping that involve predicting a constant output. We can therefore think of the readout matrix as acting not on the representation and observations directly, but on their demeaned partners, 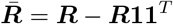 and similarly *Ō*. Then the minimal L2 readout matrix which predicts as much of the data as possible is given by the pseudoinverse:

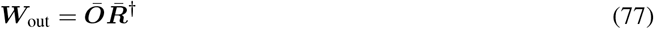

Then we can measure the reconstruction error and readout weight norm:

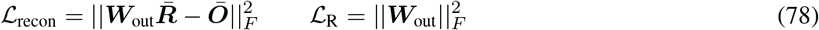

##### Recurrent Loss and Computing Constraint

The procedure for the recurrent weight is similar, except rather than mapping representation to output, we map subspaces to permuted subspaces. Hence we construct the following min-norm matrix:

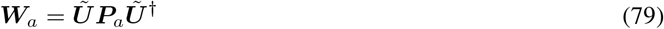

where *Ũ* corresponds to the subspace matrix without the bias (since only the non-constant parts have to be transformed by the representation), and ***P***_*a*_ implements the appropriate permutation. For example, if the task had a set of three subspaces and the action looped you through them, it would be:

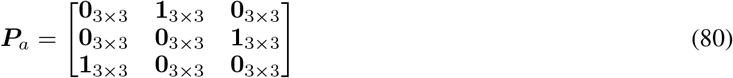

where **1**_3*×*3_ is the 3-dimensional identity matrix. Then the losses are:

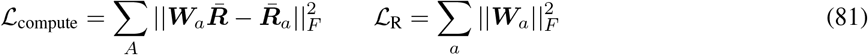

where ***R***_*a*_ is the representation after appropriate perumtation by the action. In fact we find that the computing constraint is automatically satisfied when the weight loss is included, likely because the weight losses grow enormously if the optimiser approaches a solution in which they are not, allowing us to drop it.

##### Activity

The final activity loss is simple:

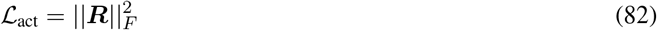

#### D.1.2 Optimising over Representation

This approach is identical to the previous one except, rather than creating ***R*** using the affine map from a set of lagged inputs, we directly optimise over ***R***. This drops the constraints that are required to ensure that the representation is only a subspace section B.1.

Otherwise the optimisation is very similar, and most of the losses are the same. One small change is that we cannot use the subspace permutation matrix above. Instead, we create the best recurrent weight matrix as:

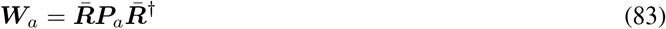

where ***P***_*a*_ implements the appropriate permutation. For example, if the task had a set of three states and the action looped you through them, it would be:

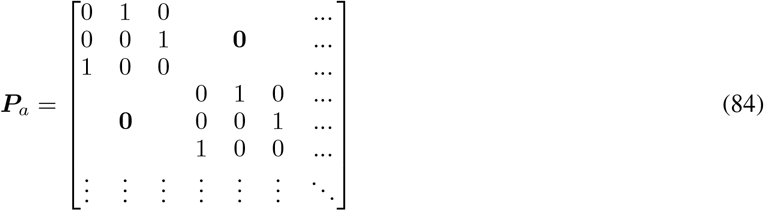

#### D.1.3 Optimising a Gated Linear RNN

The final alternative is more similar to a neural network. We specify a set of inputs, a set of outputs, and a set of actions which determine which recurrent weight matrix to use. We then construct the representation by feeding inputs and implementing the gated-linear RNN equation:

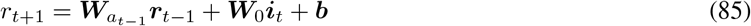

Then we follow the same procedure above, except using the explicitly parameterised ***W***_*a*_ matrices, and ignoring the computing constraint, since the RNN formulation enforces it architecturally.

### D.2 Task Details

We now outline each of the tasks. Specifying a task involves specifying the sets of memories, and the actions. Additionally, in the RNN setting, we have to specify the inputting scheme.

#### D.2.1 Subspace Parametersiation

##### Xie

On the first three timesteps we sample three observations, encoded as one-hots, then there is a delay period, before the three stimuli are recalled in order. There is a single forward action relating these, hence a conveyor belt of 5 subspaces (though see later for other action parametersiations that produce other subspace structures). We use this for fig. 4F and fig. 5E.

#### D.2.2 Neural Parameterisations

##### El-Gaby

4 observations sampled from a potential 9 without replacement and encoded one-hot, with a single forward action. Used to produce fig. 4L and fig. 5C.

##### Mushiake

3 observations sampled from a potential 4, encoded one-hot, with a single forward action. We used only the 24 sequences used in the experiment. Used to create fig. 4I.

##### Overlapping Subspace

For fig. 12, we use two sequences length 3 sequences, with a single forward action. In the deterministic case all 6 stimuli are unique. In the slightly overlapping case one stimuli overlaps between the two.

##### Contextual vs. Compositional

Sequences of length 2, each element sampled from one of three, encoded one hot, and with a single forward action. We either use only two sequences, (0, 1) and (0, 2), or we sample all possible combinations of pairs. This generates fig. 4C.

##### Panichello & Buschman

We sample 2 observations from a set of 4 either with or without replacement. We then setup a task with three timesteps: a delay period, and two potential readout futures, up or down. To create this, we setup a representation matrix of size (number of neurons) × (number of tasks × 3). The first (number of tasks) representations correspond to the delay period, the next to the recall of the top memory, the last to the bottom memory. There are two action matrices, a top and a bottom, which are constructed to move the delay period activities to their corresponding top or bottom readout. Finally, a readout matrix decodes the labels from the readout representations, either the identity of the bottom cue from the last (number of task) representations, or the top for the middle (number of task) representations. We use this for the theory parts of fig. 4D and fig. 5F.

##### Stroud

This task has two one-hot encoded stimuli and a variable delay time, between 2 steps and 4, after which there is a 2 step readout time period. There is one forward action that operates at all timepoints except when the ‘go’ cue arrives. The three tasks differ in their readout requirements. In the first, ‘cue-delay’, the readout has to predict the correct memory from the beginning of the delay period until the end of the task. In the second, ‘after-go-time’, the readout only has to predict the stimuli in the readout period. In the third, ‘just-in-time’, the representation must correctly predict from the end of the shortest delay period until the end of the task. We use this for fig. 5G.

#### D.2.3 RNN Parameterisations

##### Xie—Different Algorithms

For fig. 5I, we train gated linear RNNs on a task with up to 6 timesteps: up to 2 inputs, either 0 or 1 delay steps, and up to 2 outputs. Each sequence comprises 1 or 2 stimuli sampled from four points on a circle. We sample length two sequences without replacement, and 1 item sequences can arrive in either the first or second timestep. The actions are chosen according to two different schemes. In both cases, in order to construct an input gating network, two distinct recurrent matrices are used for the two input timesteps. Then there is a delay action. Finally there are either two distinct output actions applied after a go cue, to generate an output gating mechanism, or a single output action, to generate a conveyor belt.

### D.3 Analysis

Here we outline how we extracted various properties of the representation.

#### Simple: tuning curves, and dot-products

Figure 4C is just tuning curves, while the bottom row offig. 5G is made by using activity from the longest delay period and taking the dot-product between the activity at different time points. Further, in the case of subspace reparametrisation extracting subspace characteristics is simple, since we have direct access to the true subspace matrices, we then compute the size and alignment metrics discussed in appendix E, producing fig. 4F and fig. 5E.

#### Regression Weights

To construct neural regression weights, as in fig. 4I,L, we set up a set of regressors and run lasso regression between neural activity and the regressors.

#### Subspace properties—Neural Parametrisation

When parametrising the optimisation using the neural representation we have to first extract the subspaces. We do this using the techniques discussed in appendix E, and then construct the discussed metrics, to produce fig. 4D (theory parts), fig. 5E, and F (theory parts). To align Figure 4C with the data plot, we take advantage of the one-hot nature of the representation fig. 4L. We calculate an encoding strength for each location-lag combination by first finding each neuron tuned to that location-lag, then summing the average activity of those neurons at that location-lag. Then we average that over locations to get a subspace size measure.

#### Subspace properties—Different Algorithm Gated Linear Networks

To extract subspace properties from the gated-linear RNN network, fig. 5I, we simply pass a single observation through the network and observe its passage after different combinations of actions. We then do this for all single observations, and, for a given action sequence, we construct the subspace using the top 2 PCs of these activities. We then project each stimulus encoding into this 2D space. To measure the size of the space we simply take the average norm of these vectors. To create the information flow plots, fig. 13, we use the same subspaces, but project activity onto it from all 2 sequence trials.

## E Neural Data Analysis—Extracting (Correlated) Memory Subspaces from Data

We analysed neural data from Panichello and Buschman and Xie et al., and we are grateful to both groups for publicly sharing their data.

In both cases we extracted the sizes and alignments of the subspaces encoding each memory during the delay period of the task, and we performed similar analysis on some of our theoretical representations. In this appendix we consider methods for extracting vectors that describe the subspaces, and we construct a signed measure of alignment.

The basic problem goes as follows. We assume we have some data linearly generated by a set of variables:

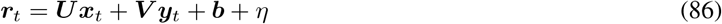

How do we estimate ***U*** and ***V*** from a dataset of {***r***_*t*_, ***x***_*t*_, ***y***_*t*_} under the noise *η*. So far, so easy. But crucially, how do we do it in a way that our estimate of the alignment of the two subspaces, ***U*** ^*T*^ ***V***, is unbiased?

The first method, averaging, is the simplest, but only works for independently sampled memories. The second, regularised regression, was used by Xie et al. and works, but is sometimes biased (not too badly). The third we invented for specifically this purpose, and works by extracting carefully chosen differences of representations that live in only one subspace avoiding interference. The fourth uses a standard debiasing approach for linear regression. We find that no method is perfect, but that each has its merits, and make conclusions based on a judicious mixture of approaches.

This section begins by outlining the preprocessing steps we performed to get the measured representations to the same state as our theoretical ones. Then, we discuss method for estimating the alignment between a set of subspaces, and problems with them, leading to our own metric. Finally, we try and estimate subspaces from data. We begin with the averaging method, how we used it to analyse the Panichello and Buschman data, and how it fails on correlated memories, producing a biased estimate. Then we describe the other methods, their results on synthetic data, and how we used them to analyse the data of Xie et al.

### E.1 Preprocessing

#### Panichello & Buschman

We worked with neural data labelled as ‘frontal’. We binned the stimuli (really a continuous angle) into 4 bins and only analysed trials on which the monkey was within half a radian of the target. We extracted the firing rate of the neurons during the last half second of the delay period.

##### Xie et al

We used all neurons from correct trials, and extracted 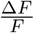 from the last second of the delay period.

### E.2 Alignment & Size Metric

Here we define our size and alignment metrics, next we will explain why we use them.

Given two subspaces, estimated by whichever method, ***U***, ***V*** ∈ ℝ^*N×K*^, we estimate their sizes as the average euclidean norm:

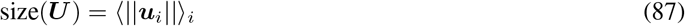

where ***u***_*i*_ is the *i*th column of ***U*** . Then the alignment is the average cosine similarity between paired vectors in each subspace:

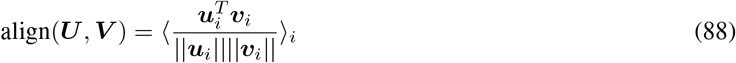

Existing alignment metrics use angles. Panichello and Buschman focus on the activity in the top three principal components, create two 2-dimensional subspaces within that space, then measure the angle between the normal vectors of the two subspaces. Alternatively, Xie et al. report the first principal angle between the extracted subspaces. Both have their shortcomings. The first method relies on projecting to 3D rather than dealing with the high-dimensional space directly. The second, by relying on only the first principal angle, introduces a bias. The principal angles are ranked, the first is smaller than the second etc. This means, even if a 2D subspace is oriented at a constant angle to another, if there is a small amount of noise it will tend to push the first principal angle smaller and the second larger, as in the synthetic data example in fig. 7. This particular effect can be removed by averaging the principal angles.

**Figure 7.**
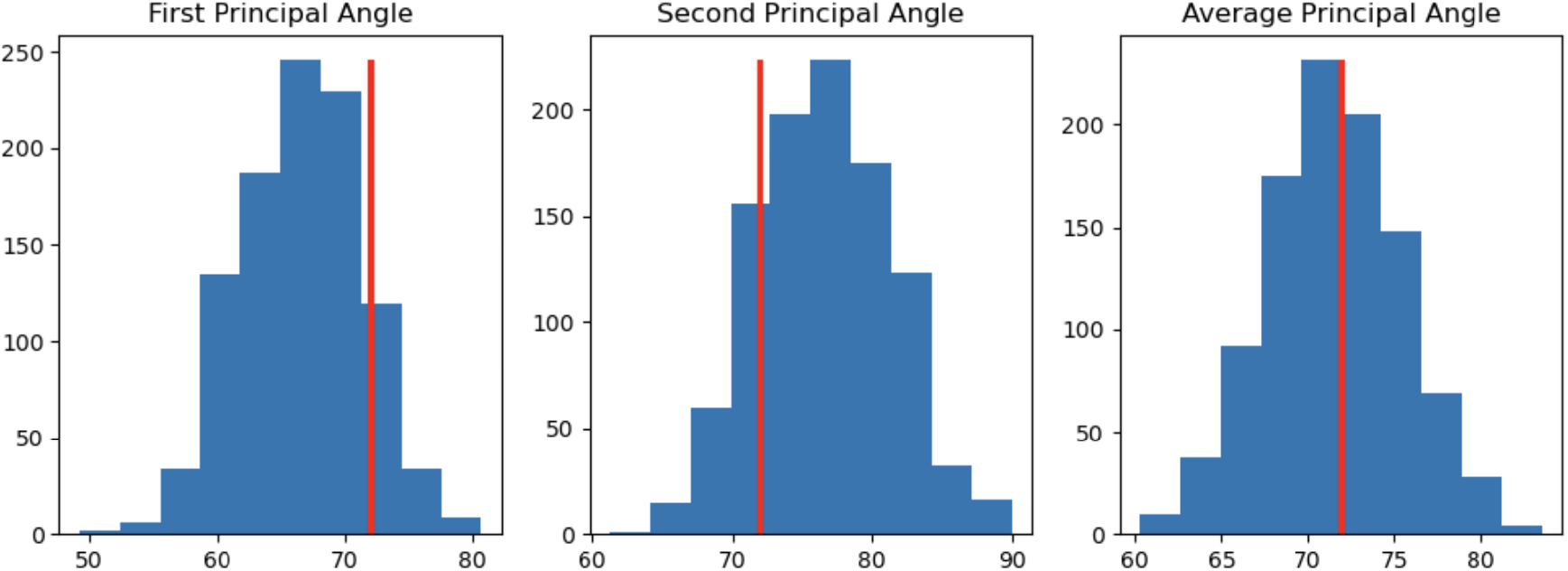
We create some fake data of two 2D subspaces that are aligned at 72^◦^, i.e. both of the two principal angles are 72^◦^, the red line. We then add a small amount of noise 1000 times and calculate the principal angles between the noisy vectors (all the vectors are length one, and we add a zero-mean gaussian noise matrix with variance 0.1). The estimate of the first principal angle is biased down, the second up, but the average appears unbiased.

However, a second problem remains. Angles are capped at 90^◦^. Let’s say your the true average principal angle is close to orthogonal. You try to estimate this from some noisy data. Perhaps some principal angles go down and some of them up. Due to the range capping the ones that went up could only go up to 90^◦^, and if they are noisy enough they might go from being slightly aligned in one direction to slightly aligned in the opposite direction. Since the values we are calculating don’t take account of the directionality of the alignment, this means that when you add noise the estimate of the principal angles slips away from orthogonal, fig 8. This suggests that any value we estimate from data is likely to be an overestimate of any close to orthogonal alignment between subspaces.

**Figure 8.**
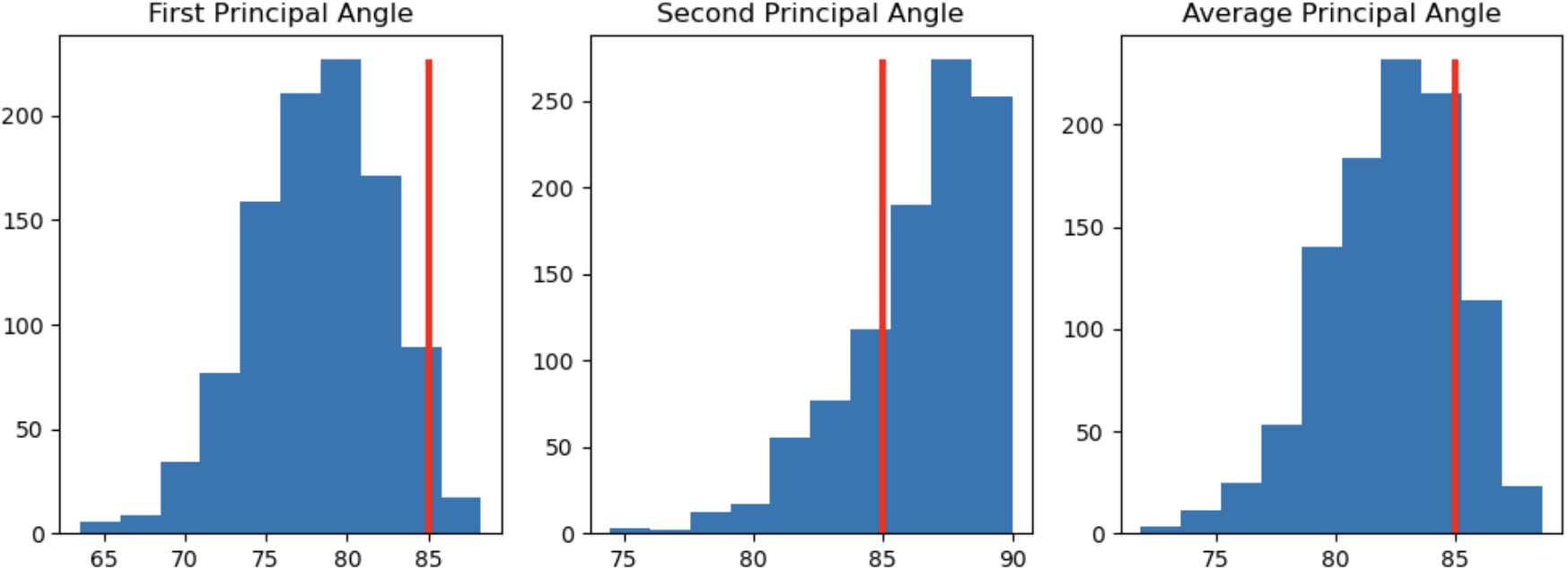
Same setting as figure 7, except now the true principal angle, the red line, is 85^◦^. We can see that all estimators, including the average principal angle are shifted downwards. This problem occurs because of the bounding of the angle. A better estimate would be a signed estimate of alignment that was 0 if the subspaces were orthogonal, positive if the encoding of the same stimulus in different subspaces align, and negative if they don’t. We therefore calculate the average cosine similarity between pairs of the same stimulus across the two subspaces, which avoids all these problems.

### E.3 Estimating Subspaces by Averaging

We begin with our first way of estimating the subspaces. If ***x***_*t*_ and ***y***_*t*_ are independent then life is easy. Averaging all ***r***_*t*_ for a given ***x*** produces an estimate of a shifted projection of ***U***, since the average of ***y***_*t*_ conditioned on ***x***_*t*_ is the same as the unconditioned average of ***y***_*t*_:

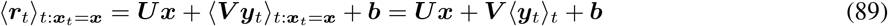

Doing this for all ***x*** creates a set of vectors, then performing PCA we create a 2D subspace that encodes these memories. To get a size estimate we project each average ***x***_*t*_ encoding onto this subspace and measure the average size of the projection, and to we calculate alignments using the previously measured metric.

#### Failing for Correlated Memories

Imagine now that you have correlated the memories (which wasn’t actually a problem for Panichello and Buschman). For example, imagine two perfectly orthogonal subspaces encoding a set of 4 direction, N, S, E, W, fig. 9. You now take the average conditioned on the first stimulus being north. Clearly the conditioned average activity in the first subspace points in a consistent direction, so far so good. However, because of the without replacement sampling, the north stimuli is never in the second subspace during these trials. Hence, in the second subspace the conditioned average points south.

**Figure 9.**
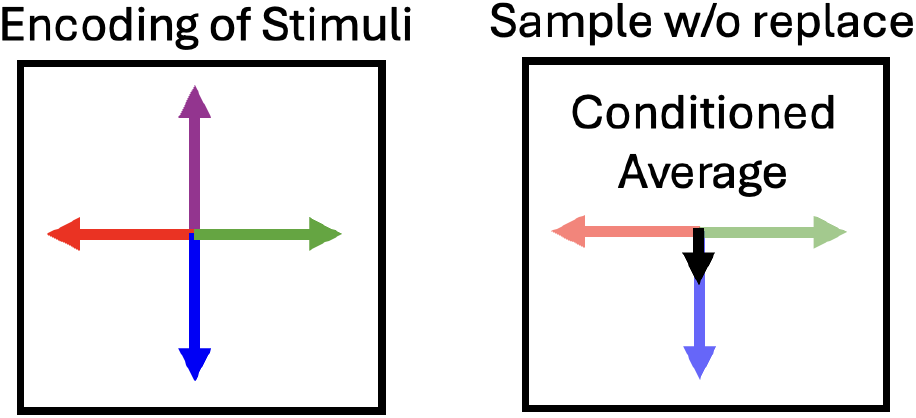
Imagine each subspace contains data pointing in the 4 cardinal directions. Between the two subspaces these directions are sampled without replacement. This means the average activity in the second subspace when the first subspace is pointing north will actually point south.

Now that you have your biased average ‘stimulus 1 = north’ activity, you take the dot product with ‘stimulus 2 = north’ average. Despite the fact the subspaces are orthogonal you find a negative alignment! The anticorrelations in the variables manifests as a predicted negative correlated between the subspaces.

### E.4 Regularised Regression

Xie et al. had to come up with a clever method to circumvent this problem: they performed regularised regression. Assume each memory is a linear combination of different subspace encodings, writing the neural activity for a sequence of two angles *θ*_1_ and *θ*_2_ as follows:

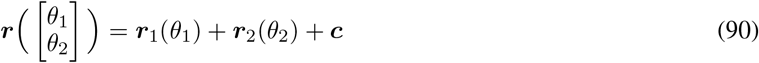

Stack each of the encoding vectors into a matrix, the coefficients of the regressors:

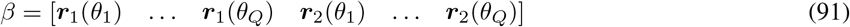

Then for each sequence you create a two-hot (in this case, because there’s two sequence elements) vector that pulls out those vectors, let’s call it 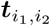. Minimise the following LASSO regression loss:

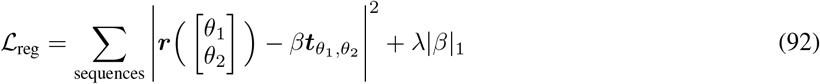

The *λ* is chosen by splitting the data in half and finding the value which causes the error on the second half of the data to be minimised. This method is much better: on noiseless data it estimates the quantity correctly, fig. 10.

**Figure 10.**
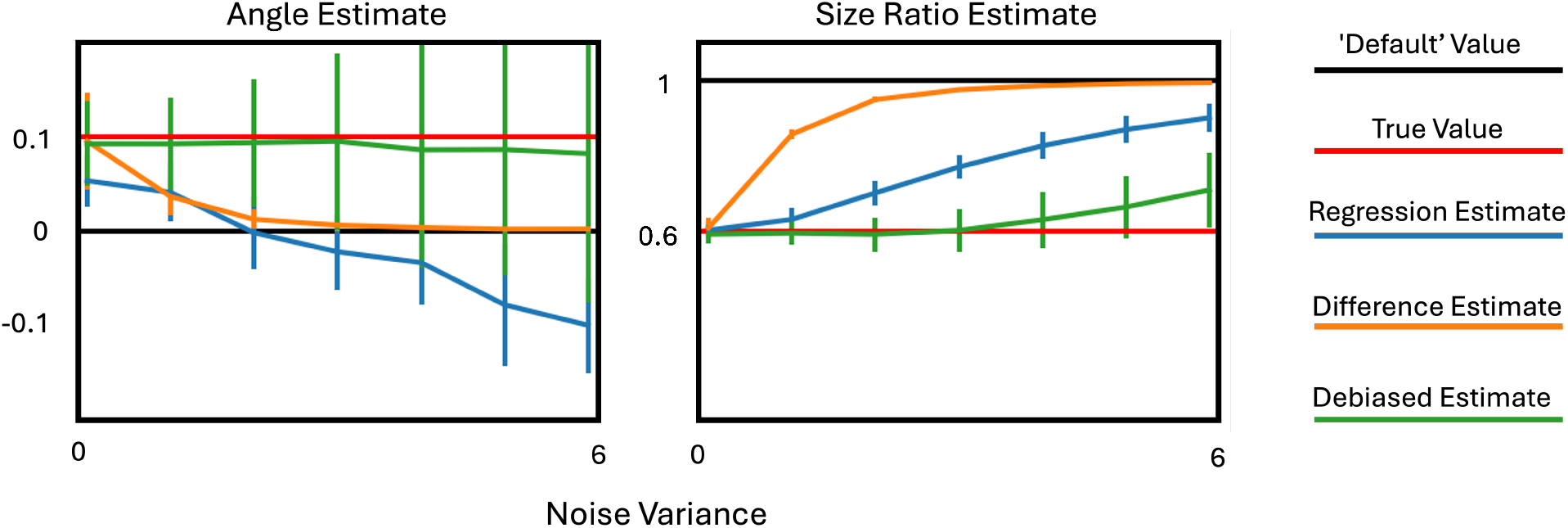
We create data from two subspaces aligned at 0.1 with size ratio 0.6 and add Gaussian noise. We use the estimators to find the size ratio and angle across 100 noise draws and plot the mean and standard deviation across different noise levels. All are good estimates at low noise levels. At high noise levels the difference method returns to ‘default’ estimates, while the angle estimated by the regression estimator heads negative. However, the regression estimator’s estimate of size is much better than the difference method’s for longer. The debiased estimator is accurate throughout, though the angle estimate has large variance. Since the debiased estimator performs best, we use it throughout.

Unfortunately, on noisy data it is biased, fig. 10. This can be more easily understood with L2 regularised regression, and the same insights appear to carry over to L1.

#### Ordinary Least Squares Estimator

The regularised L2 regression weights for mapping from the stacked matrix of regressors, ***T*** ∈ ℝ^*R×T*^, where *R* is the number of regressors and *T* the number of datapoints, to stacked neural activity ***R*** ∈ ℝ^*N×T*^ is:

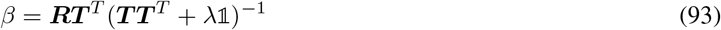

We are interested in the dot-product of the regressors. These are second order statistics, so they are governed by the covariance of the estimator:

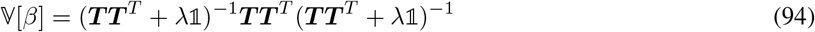

If *λ* is small (low noise), this is:

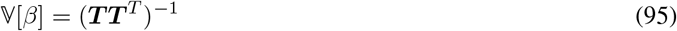

This means that, even if the subspaces are orthogonal, if the regressors are correlated (***T T*** ^*T*^ )^−1^ ≠ 0 producing aligned estimates.

Alternatively, if *λ* is large (high noise):

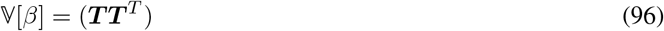

So now the alignment estimate follows the correlations of the regressors, rather than the inverse correlation matrix.

Often the elements of the correlation and inverse correlation matrix have different signs, therefore, varying *λ* will push your estimate from positively biased to negatively biased. Somewhere in the middle is a perfect *λ* value, and held-out data does quite a good job of finding it for low noise levels. However at high noise levels it fails, fig. 10. We’re not sure why, but empirically it tends to choose an overly large *λ*, leading to estimates that align with the correlations in the data.

### E.5 Difference Method

Next, we present an alternate method for estimating subspaces. We simply take the difference of encodings that differ only in the subspace we are interested in:

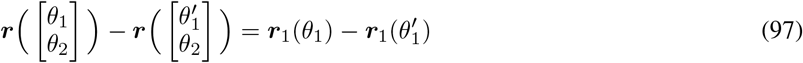

For all pairs of sequences that differ in only one element we take the difference, producing a vector that lives in one subspace (+ noise). We use the average length of these vectors as our size estimate for each subspace, and we calculate the average dot product between the same difference in two subspaces, e.g.:

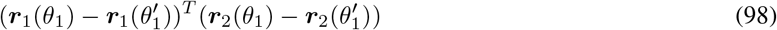

On noisy synthetic data this is also biased, fig. 10, but in ways that are conducive to our aims. As the noise increases all the alignments collapse to zero, i.e. the alignments never switch from positive to negative. This is perfect for arbitrating hypotheses like ‘these subspaces are positively aligned’. If our estimator says it is, then that is *despite* the noise, rather than aided by it, as can be the case for the regression estimator.

### E.6 De-biased Estimator

Finally, we consider a more principled approach. Given an ordinary least squares regression problem:

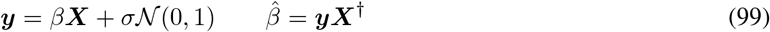

While the estimated regression weigths are unbiased, their second moments are not:

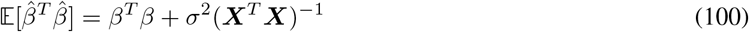

But to get an unbiased estimate we can simply remove the bias. We use the following estimator for *σ*^2^. Calling the residuals, 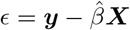:

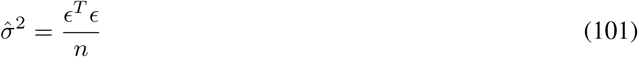

where *n* is the number of datapoints. Hence we perform OLS regression, and form the following estimator:

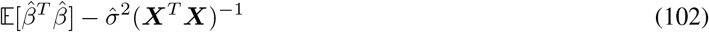

This estimator runs into one more problem, if the estimated 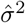 is large enough it can stop the resulting estimate from being positive semi-definite. We therefore project from our estimated covariance matrix to the positive semi-definite matrix that is closest in frobenius norm.

This estimator performs quite well on synthetic data, fig. 10, but is very noisy.

### E.7 Comparisons and Applications to Data

In fig. 11 we apply each of the three subspace estimation methods to the data of Xie et al. All three estimators find that the size of each sequence subspace decays 1 →2 →3, but to different amounts. Similarly, all three estimators find significant positive alignment between subspaces, but they differ in which subspaces are significantly aligned, and its magnitude. Given our three estimators various pros and cons, highlighted in fig. 10, how should we best combine them to draw conclusions about the data?

**Figure 11.**
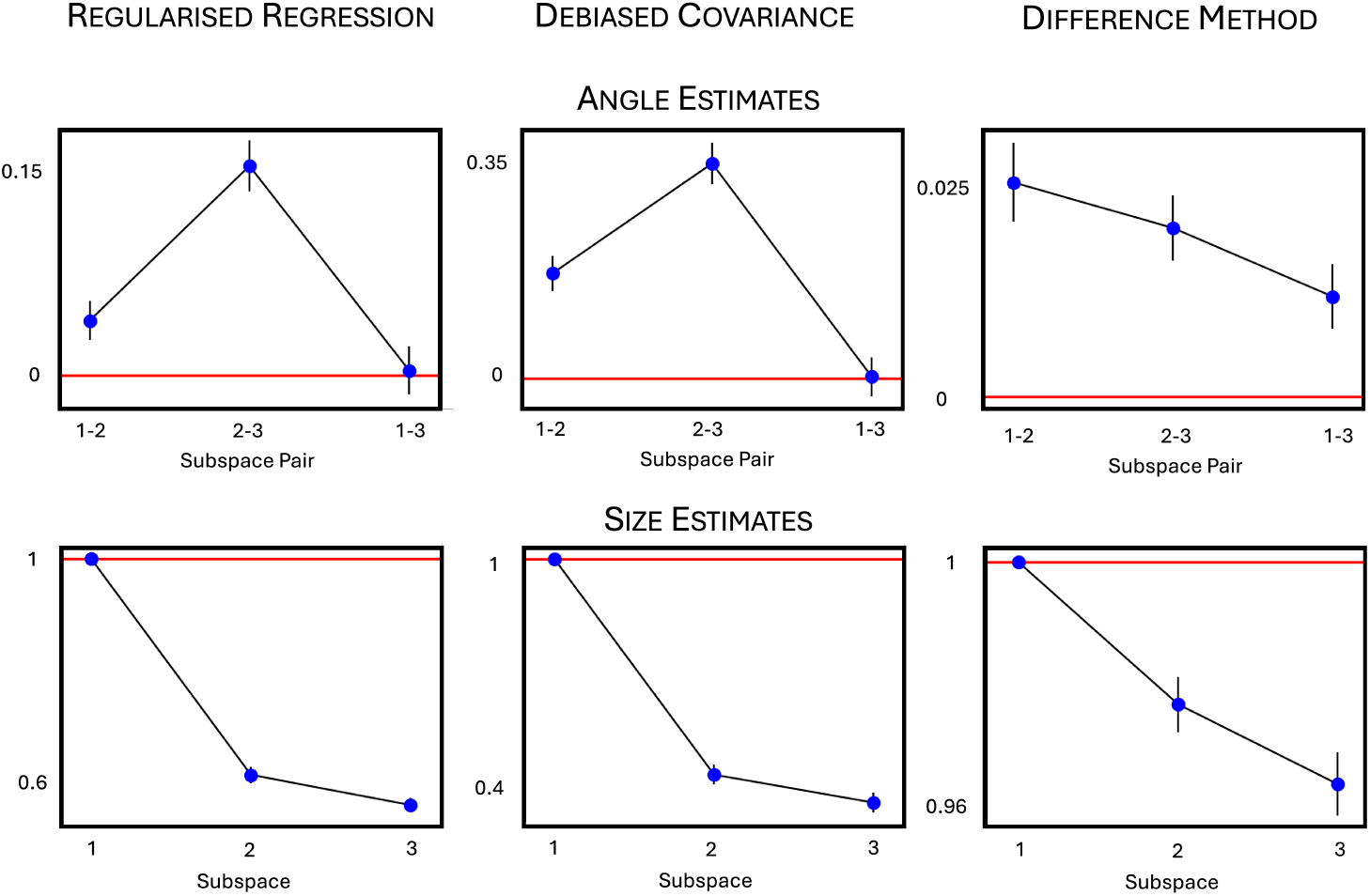
We try each of the three methods on the data from Xie et al. The top row shows the angle estimation, the bottom the size estimation. The first column uses regularised regression as in Xie et al., the second uses the debiased covariance estamator, the third uses the difference method.

We take a conservative approach. For the alignments we use the difference estimator, despite its surprisingly small values. It has the the appealing property that it never reports a subspace as being aligned when it is not— fig. 10 left, orange line. Therefore, we trust its finding that all subspaces are significantly positively aligned more than the more noisy estimates provided by either the regression method, or the (very noisy) debiased estimator. We do not make strong claims about the size of the alignments.

For the sizes, we use the debiased estimator. It appears that both the regression and the debiased estimator are relatively reliable size estimators— fig. 10 right, blue and green lines. The debiased one seems slightly more robust, though again, we don’t put huge confidence on the exact extracted magnitudes.

We therefore use this method to analyse to produce both our theory plots fig. 4D,F and fig. 5E,F, and the data plots fig. 4E and fig. 5D.

## F Supplementary Experiments and Figures

We include two supplementary figures. The first shows the limit of purely contextual coding. The second demonstrates that the representations we train learn the algorithms we suppose.

### F.1 Purely Contextual Coding

First, we consider completely dependent memories in which knowing your current stimulus can predict the rest of the sequence. Consider an agent that navigates two sequences of three elements arranged in a loop, with a single action that loops through them (blue arrow on left, fig. 12). We consider the case where there are six one-hot coded stimuli, three in each sequence, placing it in the completely dependent category. Correspondingly, in the optimal representation every neuron is ‘purely contextual’: it fires for a given stimulus, which necessarily arise in only a single sequence, fig. 12. In subspace language, the subspace encoding current and future memories are now one and the same.

**Figure 12.**
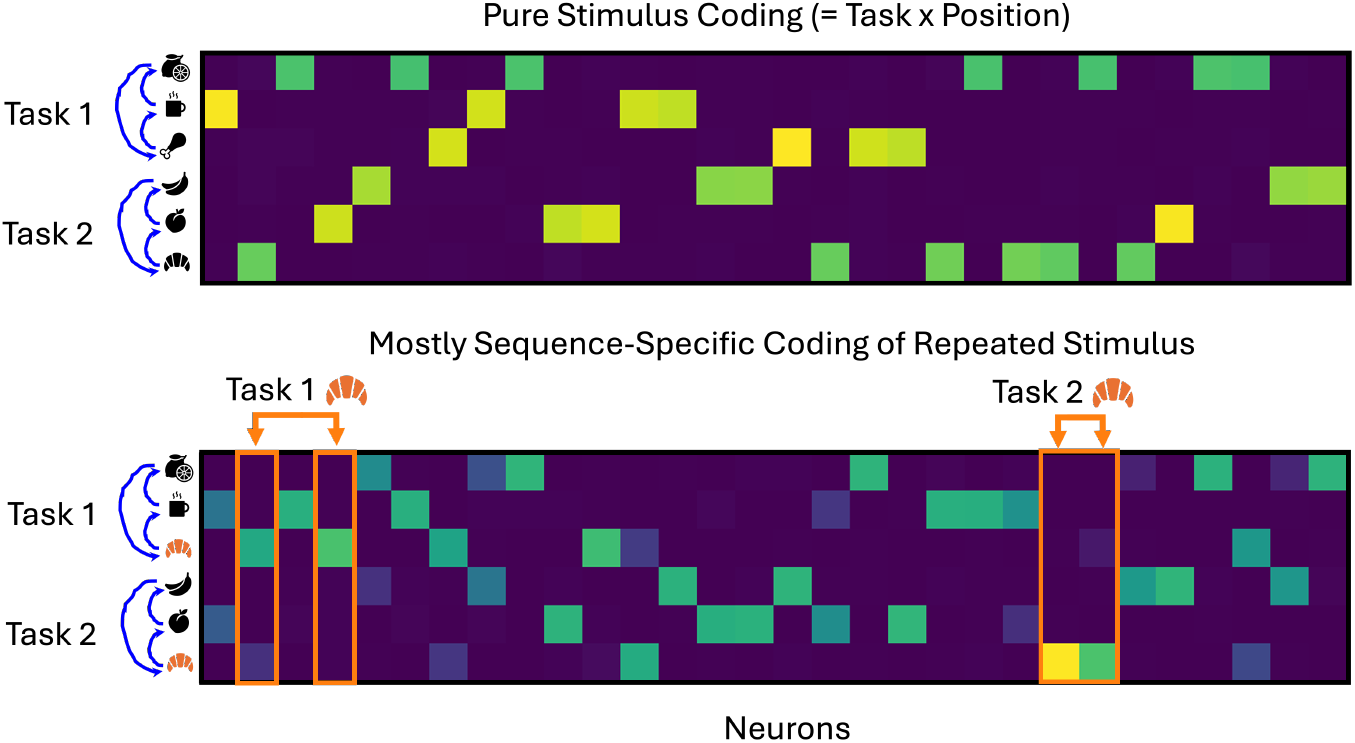
We train a representation on two length-three sequences. When there are no overlapping stimuli the optimal representation simply encodes the stimuli. Introducing one overlapping sequence element, the croissant, leads to the highlighted contextual neurons that fire at the croissant, but distinguish on the basis of which sequence.

We then introduce a minimal amount of independence, a repeated stimulus, the croissant on the bottom row of fig. 12. Since we are still very close to the determinstic end of the spectrum most neurons fire in only one sequence, and when they fire in both they have very differently sized responses. The highlighted neurons are tuned to the croissant, but only in one or other sequence, highlighting the contextual coding.

### F.2 Alternative Subspace Algorithms

Here we further discuss the subspace algorithms mentioned in section 4.3. In particular, we show the movement of the different sequence information through the network.

In fig. 13 we show the two learnt representations. Each stimulus comes from the corners of a square, and we extract the four relevant subspaces: input, storage I and II, and output. For each timestep, rows, we plot the activity in each of the subspaces. As can be seen, fig. 13B, for the gating solution each stimulus arrives, moves to its storage space, and then leaves. On the other hand, fig. 13E, the conveyor belt solution briefly sends stimulus II to the stimulus I subspace. This small difference manifests via a difference in size, fig. 13C,F, fig. 5I.

**Figure 13.**
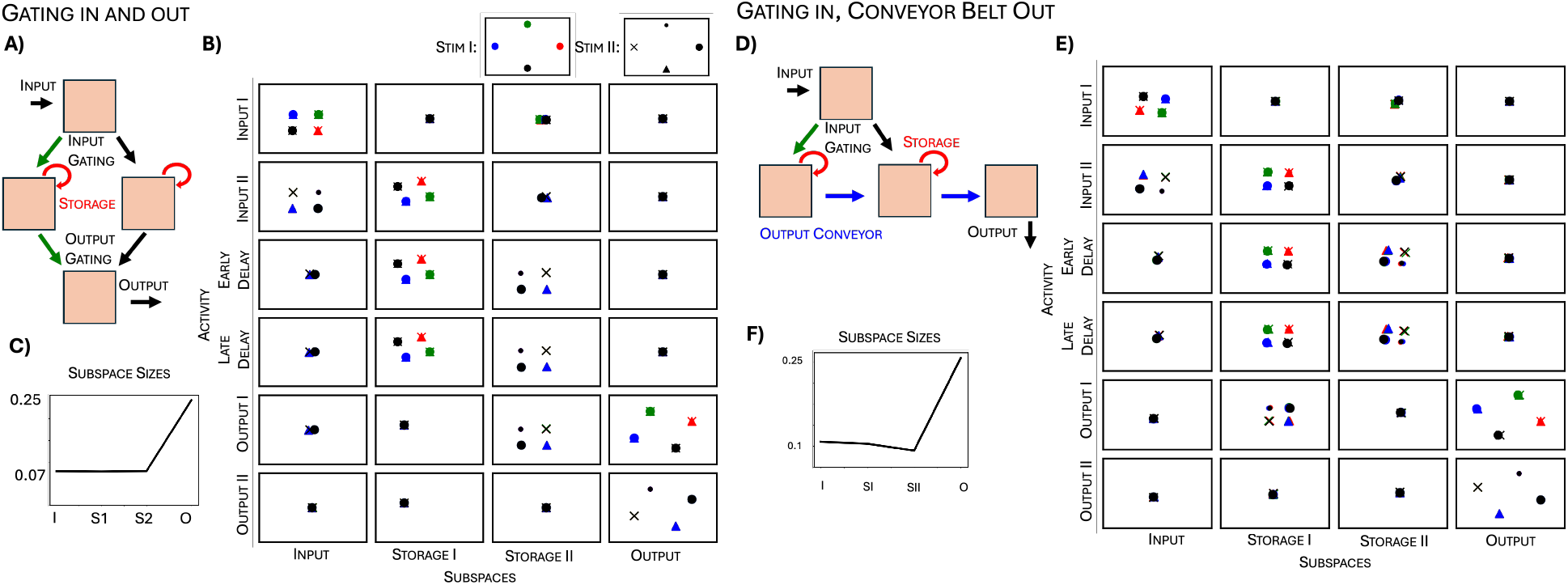
Further analysis of the subspace algorithms. **(A)** Schematic for gating solution. **(B)** Each point in these plots corresponds to the projection of the network representation onto a particular subspace. The row of the plot tells you the representation’s timestep, the colour and shape tell you the stimuli on that trial (colour = stimulus I, shape = stimulus II). Each column of plots corresponds to a different subspace. We find that the neural activity flows from input to stim I or stim II subspace, stays there, then leaves to output. **(C)** This produces subspaces in which only the output is larger. **(D-F)** Same for the network that learns to be a conveyor belt, in this case the storage II subspace is smaller than storage I, matching data.

To find these two solutions we train with multiple sequence lengths (both 2 and 1) and delays. This is because the RNN tends to take advantage of every predictability to save energy. For example, if only sequences of length 2 are used the network knows it can send stimulus I to an intermediary (smaller) subspace before it will have to be recalled. We leave further exploration of this space of solution algorithms for future work.

## Footnotes

2 This loss can also be derived as a limit of an accuracy loss with Gaussian noise on the neural activity and weights, appendix C.

3 Estimating subspace alignment from data is non-trivial. In appendix E we describe methods to do this, that we apply or reapply to existing data. The results are complex, but they confirm this discrepancy.

4 We ignore the reversing of sequences studied by Chen et al.

5 This means there need to be at least *KN*_*A*_ neurons, else the network cannot solve the task

6 The code shows it is also neither log-convex nor log-log-convex.

7 if two variables are statistically independent then knowing the value of one variable doesn’t change the distribution of the other. Range independence means knowing the value of one variable doesn’t change the range of possible values, or the support, of the other variable.

8 In fact, only a much looser property of extreme-point independence (across the set of datapoints when one variable is at its maximum or minimum value the other variables take both their maximum and minimum value) is necessary for this proof to work.

